# Peripheral sensory neuron CB2 cannabinoid receptors are necessary for both CB2-mediated antinociceptive efficacy and sparing of morphine tolerance in a mouse model of neuropathic pain

**DOI:** 10.1101/2022.05.16.492135

**Authors:** Lawrence M. Carey, Zhili Xu, Gabriela Rajic, Alexandros Makriyannis, Julian Romero, Cecilia Hillard, Ken Mackie, Andrea G. Hohmann

## Abstract

Painful peripheral neuropathy is the most common neurological complication associated with human immune deficiency virus (HIV) infection. Currently available treatments fail to provide adequate symptom relief, indicating the need for novel treatment strategies. To address this gap in knowledge, we characterized the impact of cannabinoid CB2 agonists, which lack psychoactivity associated with central CB1 activation, on antiretroviral-induced neuropathic nociception and identified cell types expressing CB2 that mediate the antinociceptive efficacy of CB2 agonists. Two structurally distinct CB2 agonists (AM1710 and LY2828360) alleviated antiretroviral-induced neuropathic pain, benefits which were absent in CB2 knockout mice. Conditional deletion of CB2 from peripheral sensory neurons eliminated the antinociceptive efficacy of CB2 agonists. We also asked whether LY2828360 treatment could reverse established morphine tolerance in the ddC-induced neuropathy model and whether CB2 expression on peripheral sensory neurons is necessary for sparing of morphine tolerance by LY2828360. The present studies suggest that CB2 activation may alleviate HIV-associated antiretroviral neuropathy and identify a previously unreported mechanism through which CB2 activation produces antinociceptive efficacy. Our results also provide the first evidence that a CB2 agonist can reverse established morphine tolerance and demonstrate that CB2 localized to peripheral sensory neurons mediates the opioid tolerance sparing efficacy of CB2 agonists.

## Introduction

In humans living with human immunodeficiency virus (HIV) infection, the development of a painful peripheral neuropathy is the most common neurological complication, with prevalence rates ranging from 20-90% (1). HIV-associated distal sensory polyneuropathy is thought to arise from two primary causes, neuropathic pain associated with toxic viral protein byproducts secreted during viral replication, or antiretroviral toxic neuropathy (ATN) caused by drugs used to combat HIV infection. Currently available pharmacotherapies used for the management of HIV-associated sensory neuropathy fail to provide adequate symptom relief (2), indicating a need for novel treatment strategies in managing HIV-associated painful neuropathy.

The endocannabinoid system has emerged as a therapeutic target for the management of HIV-associated sensory neuropathy, with several reports suggesting that cannabis alleviates pain symptoms in HIV-affected individuals (3–5). Components of cannabis which target cannabinoid type 1 receptors (CB1), such as Δ^9^-tetrahydrocannabinol (THC), promote pain relief but also produce problematic side effects (6). Cannabinoid type 2 receptor (CB2) agonists alleviate various types of pathological pain without producing unwanted side effects associated with CB1 receptor activation (7, 8). However, therapeutic efficacy of targeting CB2 to alleviate HIV-associated neuropathic pain remains largely unexplored. To date, a single report has suggested that the phytocannabinoid β-caryophyllene, which acts as a CB2 agonist, among other targets, effectively alleviates antiretroviral-associated neuropathic pain (9). However, the mechanisms by which CB2 agonists produce antinociception remain poorly understood.

Identification of cell types expressing CB2 that mediate the therapeutic benefits of CB2 ligands has remained problematic given the lack of reliable antibodies for CB2 and low levels of CB2 expression in central nervous system (10–12). To circumvent CB2 antibody specificity issues, the current studies employed a recently developed CB2 reporter mouse (CB2^f/f^) to aid in identification of cell types within nociceptive circuitry which express CB2 (13, 14). In this mouse line, the entire *cnr2* coding region is flanked by *loxP* sites, allowing for cell-type specific deletion of CB2 from cells expressing cre recombinase. To examine the contribution of peripheral sensory neuronal populations to the antinociceptive efficacy of CB2 agonists, CB2 was conditionally deleted from peripheral sensory neurons (advillin^Cre/+;^CB2^f/f^ (15)). As µ opioid receptors (MOR) present on primary afferent nociceptors are responsible for the development of tolerance to the antinociceptive efficacy of morphine (16), the present studies also sought to determine whether primary sensory CB2 expression contributes to the tolerance sparing effects of CB2 agonists.

## Results

### CB2 activation alleviates ddC-evoked mechanical and cold allodynia

Anti-retroviral treatment with ddC (25 mg/kg, i.p. x 3 injections/week x 3 weeks) induced hypersensitivity to mechanical and cold stimulation; this sensitization emerged by the second day after initiation of ddC administration and was stable for at least 14 days (p<0.001 for each time point). Morphine treatment dose dependently alleviated ddC-induced mechanical and cold hypersensitivity. The lowest dose of morphine tested (1 mg/kg, i.p.) produced a partial reversal of ddC-induced mechanical (p=0.02 vs. non-ddC treated mice, Figure 2A) and cold (p<0.001 vs. non-ddC-treated mice, Figure 2B) hypersensitivity, whereas the two highest doses (3 and 10 mg/kg i.p.) completely reversed ddC-induced mechanical and cold hypersensitivity. Treatment with the CB2 agonist AM1710 alleviated ddC-induced mechanical and cold hypersensitivity. The lowest dose of AM1710 partially reversed ddC-induced mechanical hypersensitivity (1 mg/kg, i.p., p=0.015 vs. non ddC-treated mice, Figure 2C), whereas the highest doses (5 and 10 mg/kg, i.p., p<0.001 vs. ddC-vehicle treated mice, Figure 2C) completely reversed ddC-induced mechanical hypersensitivity. AM1710 produced a full reversal of ddC-induced cold hypersensitivity at all doses evaluated (1, 5 and 10 mg/kg i.p., p<0.001 for all doses vs. ddC-vehicle treated mice). Similarly, treatment with a structurally distinct CB2 agonist, LY2828360, dose-dependently alleviated ddC-induced mechanical hypersensitivity. The lowest dose of LY2828360 (0.3 mg/kg, i.p.) produced a partial reversal of ddC-induced mechanical (p=0.004 vs. non ddC-treated mice) and cold (p=0.048 vs. non-ddC treated mice) allodynia, whereas the higher doses (1 and 3 mg/kg, i.p.; Figure 1G) produced complete reversal of ddC-induced mechanical and cold hypersensitivity (p<0.001 vs. ddC-vehicle treated mice; Figure 1H).

**Figure 1.**
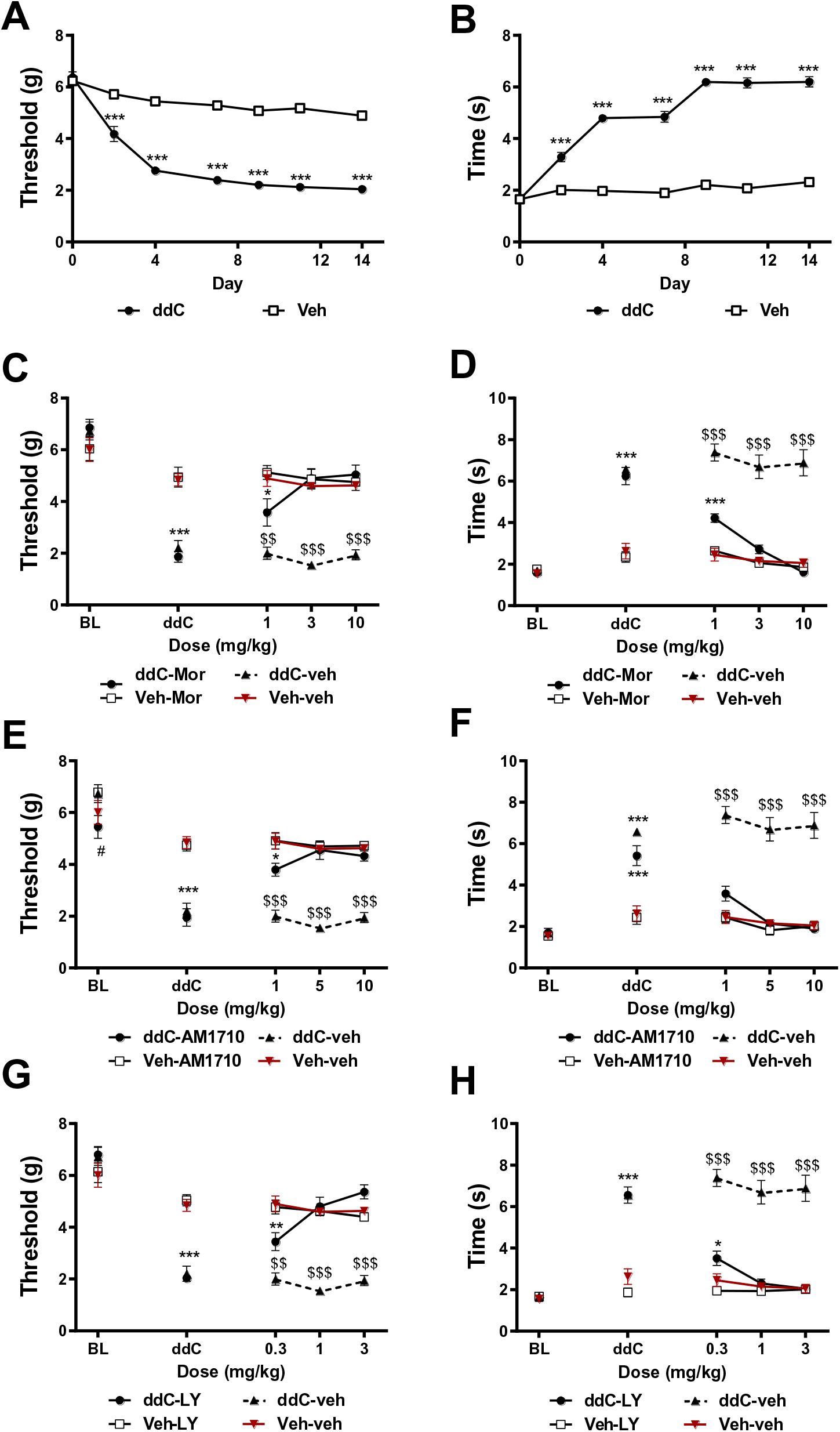
Morphine, AM1710, and LY2828360 alleviate ddC-evoked mechanical and cold allodynia. ddC induces mechanical and cold hypersensitivity. Mice treated with ddC displayed mechanical (p<0.001 for all time points, A) and cold (p<0.001 for all time points, B) allodynia on days 2-14 following onset of ddC treatments. Significant effects of ddC treatment (mechanical: F_1,44_ = 386.3, p<0.0001; cold: F_1,44_ = 541.6, p<0.0001), time (mechanical: F_6,44_ = 94.68, p<0.0001; cold: F_6,44_ = 121, p<0.0001) and ddC x time interaction (mechanical: F_6,44_ = 31.26, p<0.0001; cold: F_6,44_ = 80.17, p<0.0001) were observed. N=23 per group. Morphine (A, B), AM1710 (C, D), and LY2828360 (E, F) suppress mechanical and cold allodynia induced by ddC treatment without altering responding in mice not treated with ddC. Morphine: main effects of treatment group (mechanical: F_3,18_ = 31.59, p<0.0001; cold: F_3,18_ = 89.55, p<0.0001), dose (mechanical: F_4,18_ = 56.66, p<0.0001; cold: F_4,18_ = 80.45, p<0.0001) and treatment x dose interaction (mechanical: F_12,18_ = 11.51, p<0.0001; cold: F_12,18_ = 26.57, p<0.0001); AM1710: main effects of treatment group (mechanical: F_3,18_ = 57.32, p<0.0001; cold: F_3,18_ = 79.81, p<0.0001), dose (mechanical: F_4,18_ = 73.81, p<0.0001; cold: F_4,18_ = 60.14, p<0.0001) and treatment x dose interaction (mechanical: F_12,18_ = 11.45, p<0.0001; cold: F_12,18_ = 18.55, p<0.0001); LY2828360: main effects of treatment group (mechanical: F_3,19_ = 57.44, p<0.0001; cold: F_3,19_ = 107.9, p<0.0001), dose (mechanical: F_4,19_ = 73.71, p<0.0001; cold: F_4,19_ = 63.68, p<0.0001) and treatment x dose interaction (mechanical: F_12,19_ = 14.15, p<0.0001; cold: F_12,19_ = 26.22, p<0.0001) were observed. ***p<0.001, **p<0.01, *p<0.05 vs. non ddC-treated mice. ^$$$^p<0.001, ^$$^p<0.01 vs. all other groups. Two-way ANOVA followed by Tukey multiple comparisons test post-hoc N= 5-6 per group. BL: baseline, ddC: 2’-3’-dideoxycitidine, LY: LY02828360, Mor: morphine.

**Figure 2.**
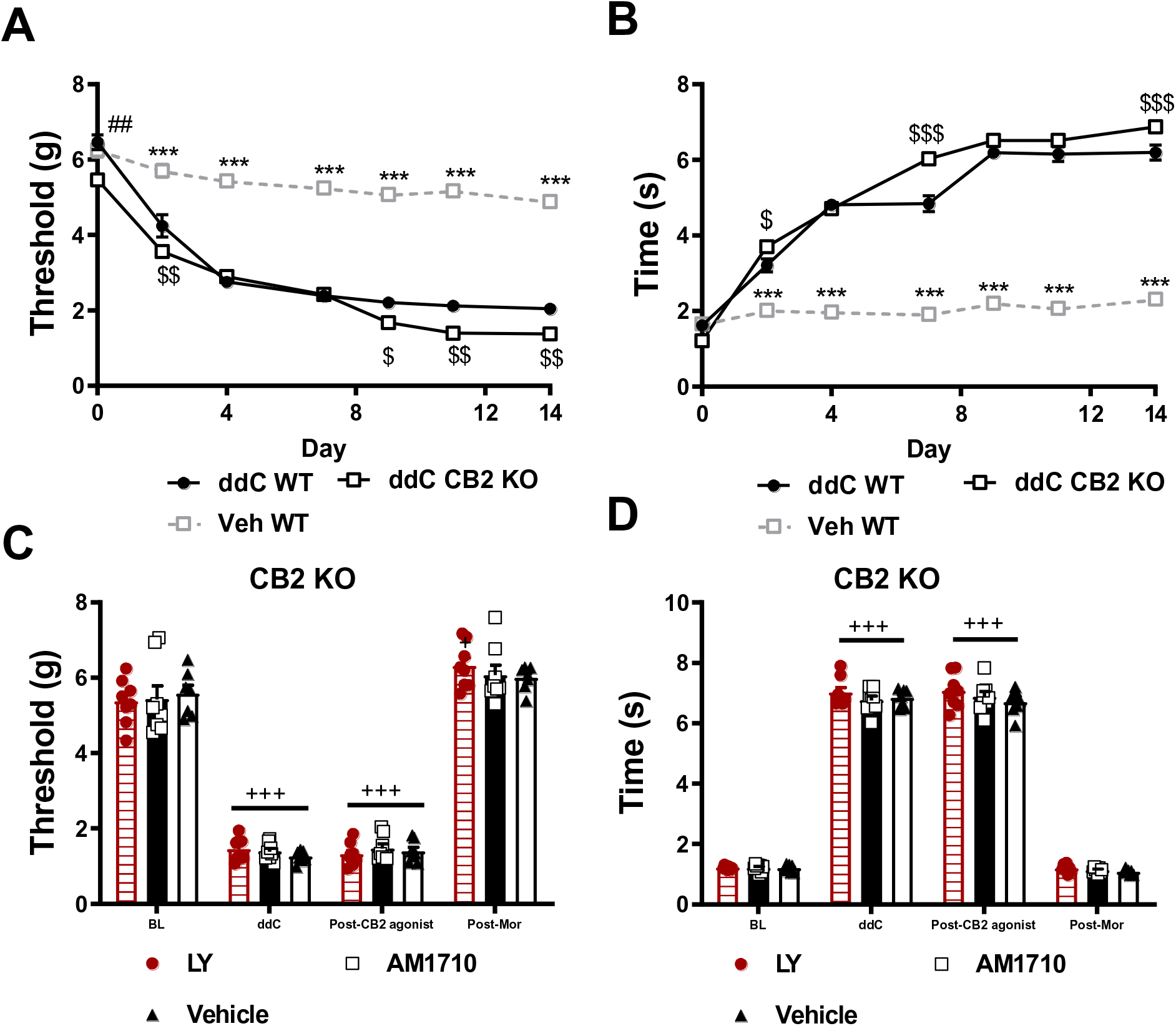
The antinociceptive effects of AM1710 and LY2828360 are absent in CB2 KO mice. Neither AM1710 nor LY2828360 alleviate ddC-evoked neuropathic nociception in CB2 KO mice. CB2 KO mice displayed slightly lower mechanical paw withdrawal thresholds at baseline relative to WT mice (p=0.0004). CB2 KO and WT mice treated with ddC displayed lower mechanical paw withdrawal thresholds on days 2-14 post initiation of ddC dosing relative to vehicle-treated mice (p<0.001 for all observations, A). CB2 KO mice displayed lower paw withdrawal thresholds relative to WT mice treated with vehicle on days 2 (p=0.003), 9 (p=0.03), 11 (p=0.002) and 14 (p=0.003). CB2 KO and WT mice treated with ddC display elevated levels of cold hypersensitivity from day 2-14 following onset of ddC treatments (p<0.001 for all observations, B). CB2 KO mice display elevated levels of cold hypersensitivity on days 2 (p=0.02), 7 (p<0.001) and 14 (p<0.001) following onset of ddC treatments. Main effects of ddC (mechanical: F_2,66_ = 439.8, p<0.0001; cold: F_6,66_ = 480.4, p<0.0001), time (mechanical: F_6,66_ = 213.8, p<0.0001; cold: F_6,66_ = 418.5, p<0.0001) and ddC x time interactions (mechanical: F_12,66_ = 22.75, p<0.0001; cold: F _12,66_ = 83.78, p<0.0001) were observed. ^##^p<0.01 vs. CB2 KO, ^$$$^p<0.001, ^$$^p<0.01, ^$^p<0.05 vs. WT ddC, ***p<0.001 vs. all other groups. Two-way ANOVA followed by Tukey multiple comparisons test post-hoc. N= 23 per group. WT: wild type, CB2 KO: CB2 knockout, ddC: 2’-3’-dideoxycitidine. LY2828360 (3 mg/kg, i.p.) and AM1710 (10 mg/kg, i.p.) fail to alleviate ddC-induced mechanical (C) and cold allodynia (D) in CB2 KO mice, whereas morphine (10 mg/kg, i.p.) restores mechanical (C) and cold (D) allodynia to pre-ddC levels in the same mice. Main effects of treatment in LY2828360-treated mice (mechanical: F_3,7_=252.5, p<0.0001; cold: F_3,7_=768.9, p<0.0001), AM1710-treated mice (mechanical: F_3,7_=109.1, p<0.0001; cold: F_3,7_=814.4) and vehicle-treated mice (mechanical: F_3,6_=297.3, p<0.0001; cold: F_3,6_=967.8, p<0.0001). ^+++^p<0.001, ^+^p<0.05 vs. baseline. One-way repeated measures ANOVA, Tukey multiple comparisons test post-hoc. N= 7-8 per group. BL: baseline, ddC: 2’-3’-dideoxycitidine, LY: LY2828360, Mor: morphine.

### AM1710 and LY2828360 fail to alleviate ddC-evoked mechanical and cold allodynia in mice lacking CB2 receptors

CB2 knockout mice (KO) developed ddC-induced mechanical and cold hypersensitivity at levels largely equivalent to wild type (WT) mice. At baseline, CB2 KO mice displayed modestly lower paw withdrawal thresholds relative to WT mice treated with ddC and vehicle (p<0.01; Figure 2A). ddC reduced mechanical paw withdrawal thresholds in both genotypes on days 2-14 (p<0.001 for each time point) following initiation of ddC dosing relative to WT mice treated with vehicle (Figure 2A), consistent with development of mechanical allodynia. CB2 KO mice displayed lower withdrawal thresholds than WT mice treated with ddC at days 2 (p=0.003), 9 (p=0.03), 11 (p=0.002) and 14 (p=0.003) following initiation of ddC dosing (Figure 2A). ddC also increased acetone-evoked nocifensive behavior in both genotypes on days 2-14 (p<0.001 for each time point; Figure 2B). However, CB2 KO mice displayed elevated levels of cold hypersensitivity on days 2 (p=0.02), 7 (p<0.001) and 14 (p<0.001) relative to WT mice treated with ddC. ddC induced mechanical and cold allodynia in all CB2 KO groups (p<0.0001 for all treatment groups). Most relevant to our studies, in CB2 KO mice neither LY2828360 (3 mg/kg, i.p. mechanical: p=0.85, cold: p=0.99) nor AM1710 (10 mg/kg, i.p. mechanical: p=0.93; cold: p=0.96) altered mechanical or cold responding relative to vehicle (mechanical: p=0.81; cold: p=0.92) treatment (Figure 2C and 2D). By contrast, morphine (10 mg/kg, i.p.) treatment completely reversed ddC evoked mechanical and cold allodynia in all CB2 KO groups (p<0.0001 for all groups; Figure 2C and D).

### CB2 agonists produce comparable levels of antinociception in male and female mice

Male and female mice exhibited comparable levels of ddC-evoked mechanical and cold allodynia on days 2-14 of the testing period (data not shown). All compounds tested showed a similar time course of anti-allodynic effects. AM1710 (10 mg/kg, i.p. Figure 4A,B), LY2828360 (3 mg/kg, i.p. Figure 4C,D) and morphine (10 mg/kg, i.p. Figure 4E,F) reversed ddC-evoked mechanical and cold allodynia from 0.5-4.5 hours post-administration (p<0.0001 for all comparisons (Figure 4B). In female mice, AM1710 lowered cold responsiveness 24 hours-post injection relative to vehicle (p=0.02; Figure 4B) Male mice showed slightly elevated cold responsiveness at 0.5 (p=0.007), 2.5 (p=0.04), 4.5 (p=0.04), and 24 hours (p=0.03) post vehicle treatment relative to female mice similarly treated with vehicle (Figure 4D). Male mice treated with LY2828360 displayed slightly lower cold responsiveness 24 hours post-administration (p=0.002) relative to female mice treated with vehicle (Figure 4D). Male mice treated with morphine also displayed higher mechanical paw withdrawal thresholds 24 hours post-administration relative to female mice similarly treated with morphine (p=0.04; Figure 4E). At 24 hours post-injection, female mice treated with morphine showed lower cold responsiveness relative to female mice treated with vehicle (p=0.0004; Figure 4F). AM1710, LY2828360 and morphine did not alter mechanical or cold responsiveness in the absence of ddC (data not shown).

**Figure 3.**
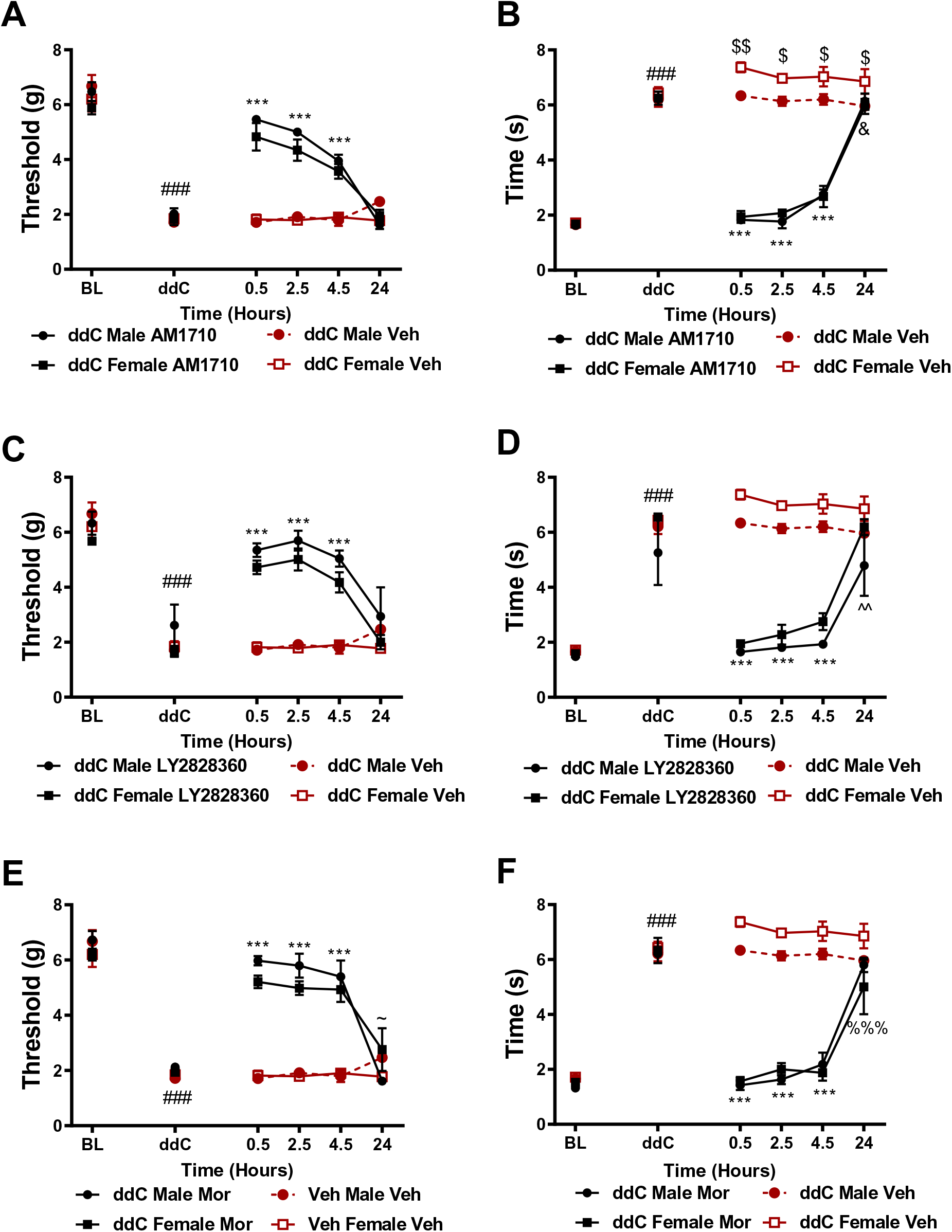
Time course and comparison of the anti-allodynic efficacy of CB2 agonists in male and female mice. AM1710 (A,B), LY2828360 (C,D) and morphine (E,F) alleviated ddC evoked mechanical and cold hypersensitivity at comparable levels in male and female mice at 0.5, 2.5 and 4.5 hours post-administration relative to ddC-treated mice receiving vehicle. Main effects of ddC for AM1710: (mechanical: F_5,60_ = 186.3, p<0.0001; cold: F_5,60_ = 296.4, p<0.0001), LY2828360: (mechanical: F_5,60_ = 77.09, p<0.0001; cold: F_5,60_ = 89.54, p<0.0001) and morphine: (mechanical: F_5,60_ = 123.4, p<0.0001; cold: F_5,60_ = 138.9, p<0.0001), treatment for AM1710: (mechanical: F_3,12_ = 38.82, p<0.0001; cold: F_3,12_ = 117, p<0.0001), LY2828360 (mechanical: F_3,12_ = 28.09, p<0.0001; cold: F_3,12_ = 46.43, p<0.0001), and morphine (mechanical: F_3,12_ = 49.93, p<0.0001; cold: F_3,12_ = 87.59, p<0.0001), and ddC x drug interaction for AM1710 (mechanical: F_15,60_ = 16.54, p<0.0001; cold: F_15,60_ = 50.94, p<0.0001), LY2828360 (mechanical: F_15,60_ = 8.735, p<0.0001; cold: F_15,60_ = 15.45, p<0.0001) and morphine (mechanical: F_15,60_ = 15.48 p<0.0001; cold: F_15,60_ = 25.93, p<0.0001). ^###^p<0.0001 all groups vs. baseline. ***p<0.0001 vs. veh-treated male and female, ^$$^p<0.01, ^$^p<0.05 vs. female veh, ^&^p<0.05 female AM1710 vs. female veh, ^∼^p<0.05 male Mor vs. female Mor, ^%%%^p<0.001 female Mor vs. female veh. Two-way ANOVA, Tukey multiple comparisons test post-hoc. (N=4 per group). Mor: morphine.

**Figure 4.**
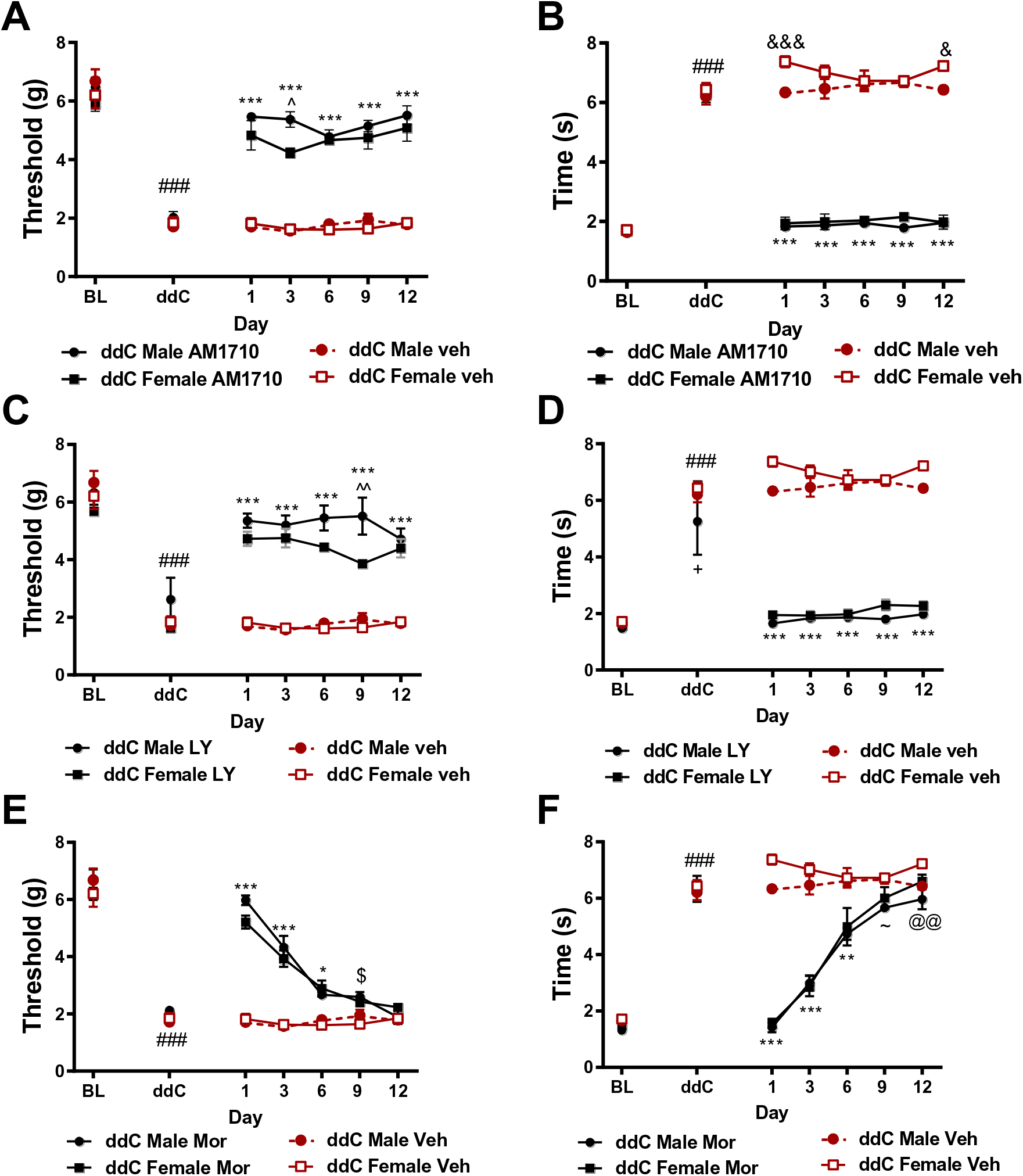
CB2 agonists display sustained antinociceptive efficacy with repeated dosing whereas opioid analgesics do not. AM1710 (A, B) and LY2828360 (C, D) produce sustained anti-allodynic efficacy following chronic administration and at comparable levels in male and female mice. Tolerance develops to the pain relieving effects of morphine (E, F). Main effects of ddC for AM1710 (mechanical: F_6,72_ = 124.6, p<0.0001; cold: (F_6,72_ = 247.3, p<0.0001), LY2828360 (mechanical: F_6,72_ = 86.43, p<0.0001; cold: F_6,72_ =89.94, p<0.0001) and morphine (mechanical: F_6,72_ = 274.6, p<0.0001; cold: F_6,72_ = 215.9, p<0.0001); treatment for AM1710 (mechanical: F_3,12_ = 108, p<0.0001; cold: F_3,12_ = 504.6, p<0.0001), LY2828360 (mechanical: F_3,12_ = 61.48, p<0.0001; cold: F_3,12_ = 407.6, p<0.0001) and morphine (mechanical: F_3,12_ =33.51, p<0.0001; cold: F_3,12_ = 61, p<0.0001); and ddC x treatment interactions for AM1710 (mechanical: F_18,72_ = 15.69, p<0.0001; cold: F_18,72_ = 62.47, p<0.0001), LY2828360 (mechanical: F_18,72_ = 11.13, p<0.0001; cold: F_18,72_ = 21.65, p<0.0001) and morphine (mechanical: F_18,72_ = 18,36, p<0.0001; cold: (F_18,72_ = 24.04, p<0.0001) were observed. ^###^p<0.0001 vs. baseline, ***p<0.0001 vs. veh-treated male and female, ^&&&^p<0.0001, ^&^p<0.05 female veh vs. male veh, ^^^^p<0.01 male LY2828360 vs. female LY2828360, ^$^p<0.05 male Mor vs. female veh, ^∼^p<0.05 male Mor vs. veh, ^@@^p<0.01 male Mor vs. female veh. Two-way ANOVA, Tukey multiple comparisons test post-hoc. (N=4 per group).

### CB2 agonists, but not morphine, display sustained anti-allodynic efficacy with chronic dosing

AM1710 (10mg/kg, i.p. A, B) and LY2828360 (3 mg/kg, i.p. C, D) produced sustained antinociceptive efficacy, at similar levels between male and female mice, alleviating ddC-induced mechanical and cold hypersensitivity throughout the 12 day chronic dosing period (p<0.0001 for all time points) relative to same sex mice treated with vehicle. In comparison, the antinociceptive effects of morphine were largely absent after 6 days of repeated injections (Figure 4E, F). (A) Male mice treated with AM1710 displayed higher mechanical paw withdrawal thresholds on day 3 of administration (p=0.01) relative to female mice treated with AM1710. (Figure 4B) Female mice treated with vehicle displayed transient increases in cold responsiveness relative to male mice similarly treated with vehicle on days 1 (p=0.0007) and 12 of injection (p=0.02). Male mice receiving LY2828360 displayed higher mechanical paw withdrawal thresholds on day 9 of chronic dosing (p=0.002) relative to female mice similarly treated with LY2828360 (Figure 4C). ddC-treated male mice displayed lower cold responsiveness relative to ddC-treated female mice receiving either vehicle or LY2828360 (p=0.02) (Figure 4D). Morphine alleviated ddC-induced mechanical hypersensitivity on day 1 (p<0.0001), 3 (p<0.0001) and 6 (p=0.03) of administration relative to vehicle treatment (Figure 4E). ddC-treated male mice treated with morphine displayed slightly higher mechanical withdrawal thresholds on day 9 of daily dosing relative to female ddC-treated mice receiving vehicle (p=0.02, Figure 4E). Morphine alleviated ddC-evoked cold hypersensitivity, at comparable rates between male and female mice, on days 1 (p<0.0001), 3 (p<0.0001) and 6 (p=0.001) of administration. Male mice showed lower cold responsiveness relative to male and female mice treated with vehicle on day 9 (p=0.03) and female mice treated with vehicle (p=0.006) on day 12 of administration. AM1710, LY2828360 and morphine did not alter mechanical withdrawal thresholds or cold responsiveness in mice treated with vehicle in lieu of ddC (data not shown).

#### Impact of AM1710, Morphine and LY on mRNA expression levels of cytokines and chemokines in lumbar spinal cord

To identify the effect of toxic challenge with ddC (25 mg/kg i.p., administered 3 times a week for 3 weeks) at the molecular level, we compared mRNA expression levels of a battery of genes in lumbar spinal cords derived from mice receiving either ddC or saline at the same times. Lumbar spinal cord mRNA expression levels of IL-1β (*p*=0.04), TNF-α (*p*=0.016) and CCL2 (i.e. MCP-1; *p*=0.047) were increased in mice that received ddC relative to vehicle treatment (Figure 5). By contrast, CNR2 (*p*=0.8768), IL-10 (*p*=0.310), CXCL12 (*p*=0.991) and CXCR4 (*p*=0.599) mRNA expression levels were not altered by ddC treatment. We further analyzed IL-1β, TNF-α, and CCL2 mRNA expression levels in lumbar spinal cords derived from ddC-treated mice chronically treated (once per day x 12 days, i.p.) with AM1710 (10 mg/kg), morphine (10 mg/kg) or LY2828360 (3 mg/kg). AM1710 decreased mRNA expression levels of IL-1β (*p*=0.057, Figure 6A), TNF-α (*p*=0.016, Figure 6B) and CCL2 (*p*=0.031, Figure 6C), cytokines and chemokines that were elevated by ddC treatment. Morphine reduced mRNA expression levels of IL-1β (*p*=0.0278, Figure 6D) and TNF-α (*p*=0.0349, Figure 6E) but not CCL2 (Figure 6F) in ddC-treated mice. LY2828360 did not alter IL-1β (*p*=0.963, Figure 6G), TNF-α (*p*=0.363, Figure 6H) or CCL2 (*p*=0.085, Figure 6I) mRNA expression levels in ddC-treated mice at the same timepoint.

**Figure 5.**
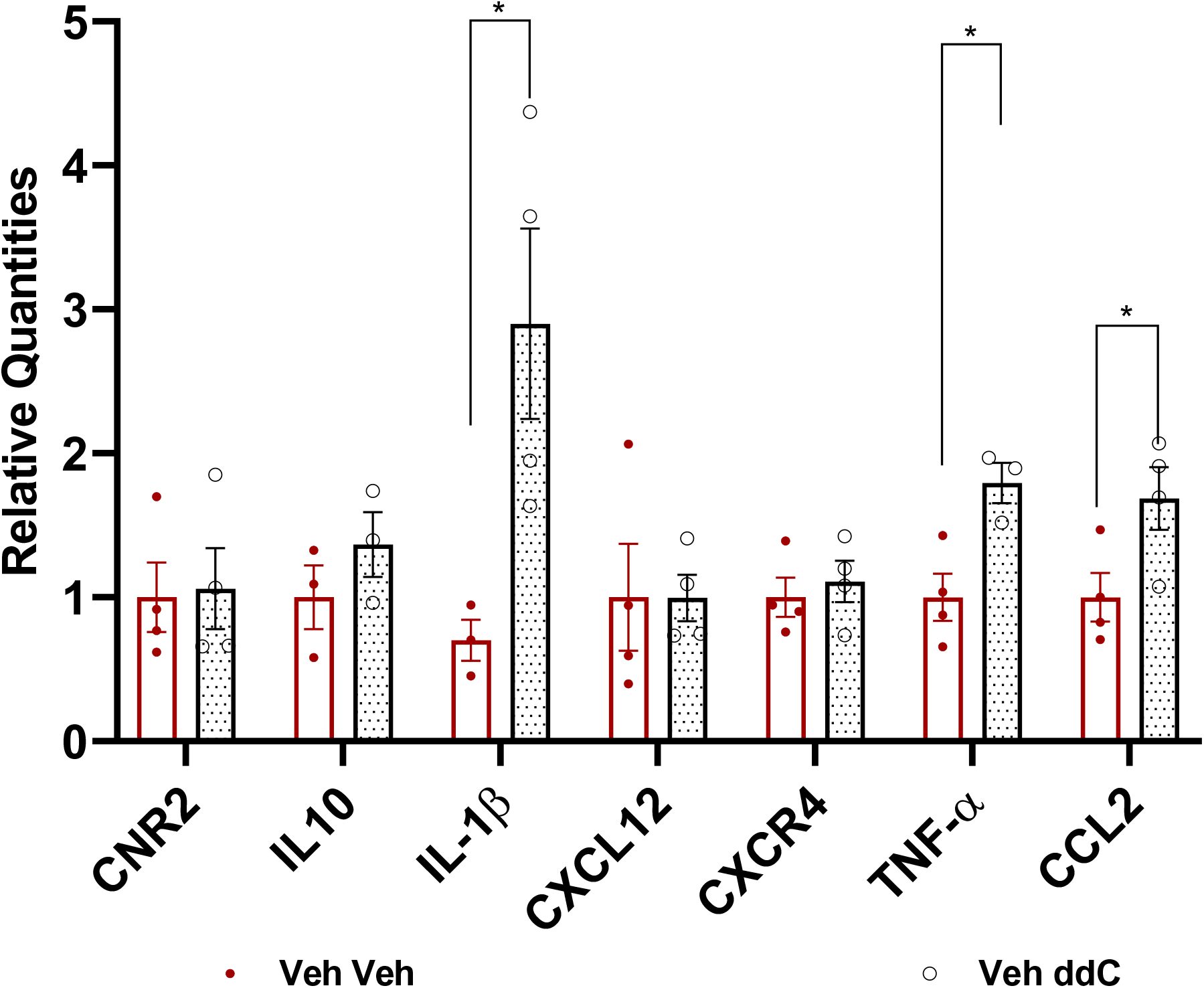
Impact of toxic challenge with ddC on CB2, cytokine and chemokine mRNA expression levels in lumbar spinal cord derived from male wildtype mice. ddC treatment produced an upregulation of IL-1β, TNF-α, and CCL2 in the lumbar spinal cord compared with the saline group similarly injected with vehicle (i.p.). Values were calculated using the 2(^-ΔΔ^Ct) method. GAPDH was used as reference gene. Data are expressed as mean ± S.E.M. (n=4 per group). * p<0.05 vs. vehicle, unpaired sample two tailed t-test.

**Figure 6.**
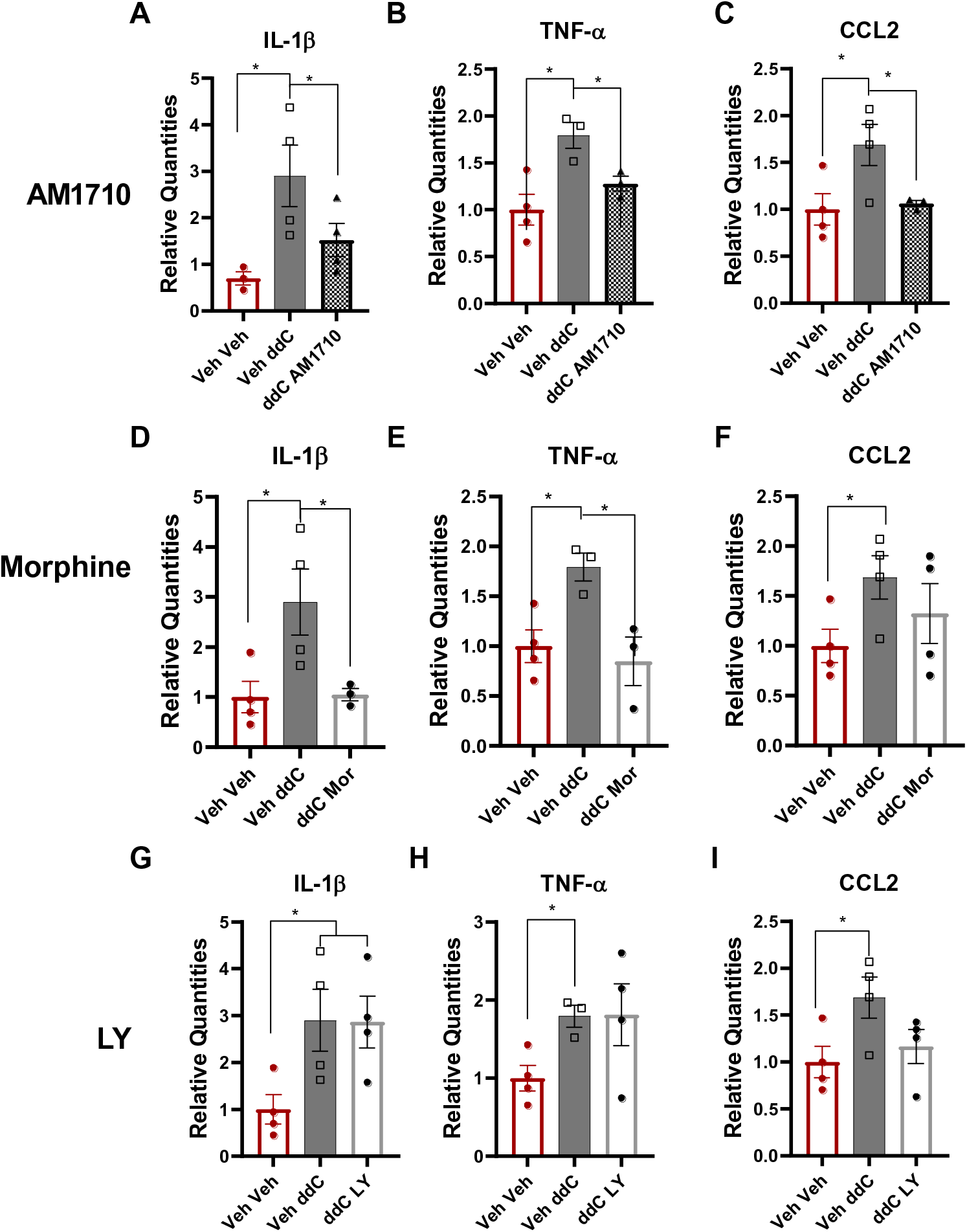
Impact of ddC and systemic (i.p.) administration of AM1710, morphine and LY2828360 on expression levels of IL-1β, TNF-α and CCL2 mRNAs in lumbar spinal cord derived from male wildtype mice. (A, B, C) AM1710 blunts ddC-evoked increases in IL-1β, TNF-α and CCL2 mRNAs. Morphine blunts ddC-induced increases in IL-1β, TNF-α mRNAs (D,E,F), whereas LY2828360 fails to alter ddC-induced increases in IL-1β, TNF-α and CCL2 mRNAs (LY; G, H, I). ddC treatment (Veh ddC) induced upregulation of IL-1β, TNF-α and CCL2 mRNAs compared to saline treated groups that were similarly injected with vehicle (Veh-Veh). Values were calculated using the 2(^-ΔΔ^Ct) method with GAPDH was used as reference gene and all groups compared to the Veh Veh group. Data are expressed as mean ± S.E.M. (n=4 per group). * p<0.05, one-way ANOVA. Mor: morphine, LY: LY2828360.

#### Immunohistochemical mapping of CB2 receptor expression in CB2^f/f^ mice

Next, we performed immunohistochemical experiments to localize CB2 protein expression throughout the brain, spinal cord, DRG, and spleen of male and female CB2^f/f^ mice sacrificed after 14 days of initiation of treatment with ddC or vehicle. Robust anti-GFP immunoreactivity was observed in the spleens of male and female CB2^f/f^ mice treated with ddC or vehicle (Figure 7C,F). Direct GFP fluorescence was below detection threshold (Figure 7B,E). GFP and anti-GFP immunoreactivity were below detection threshold in the lumbar spinal cord, brain, and lumbar L3-6 DRG of mice treated with vehicle or ddC (data not shown). Anti-GFP immunostaining was also carried out using wild type C57BL6/J mice as negative controls. As expected, anti-GFP immunoreactivity was absent in the spleens, spinal cord, and brains of WT mice processed under identical conditions (data not shown).

**Figure 7.**
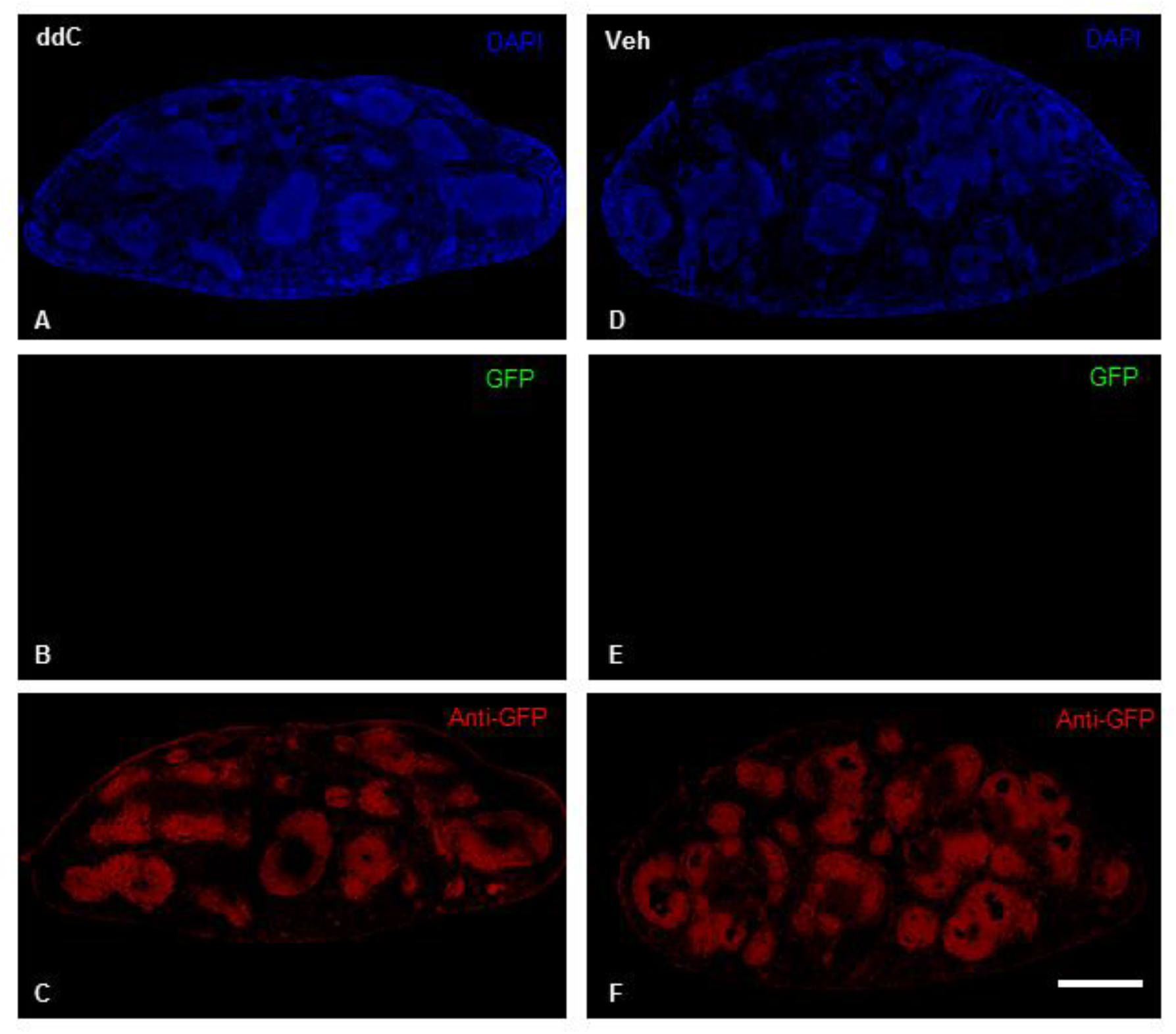
Anti-GFP-like immunoreactivity is detectable in the spleens of CB2^f/f^ mice. Robust anti-GFP-like immunoreactivity was located in the spleens of CB2^f/f^ mice treated with ddC (A, C) or vehicle (D, F). Sudan black treatment quenched native eGFP expression (B, E). Scale bar equal to 1 mm.

To circumvent possible concerns that limited sensitivity of IHC may mask low levels of protein expression and that chemical methods used to quench (auto)fluorescence in the tissues examined, we conducted qRT-PCR experiments to determine whether CB2 and GFP mRNA was present in spleen, spinal cord and DRG of CB2^f/f^ and wild type C57BL6/j mice. CB2 mRNA was detectable in the spleens, lumbar spinal cord and DRG at comparable levels in CB2^f/f^ and WT mice. Moreover, GFP mRNA was present at very high levels in the spleen, and at lower levels in both lumbar spinal cord and DRG of CB2^f/f^ mice (Figure 8D-F). However, as expected, GFP mRNA was not reliably detected in spleen, spinal cord and DRG of WT mice lacking GFP (Figure 8D-F).

**Figure 8.**
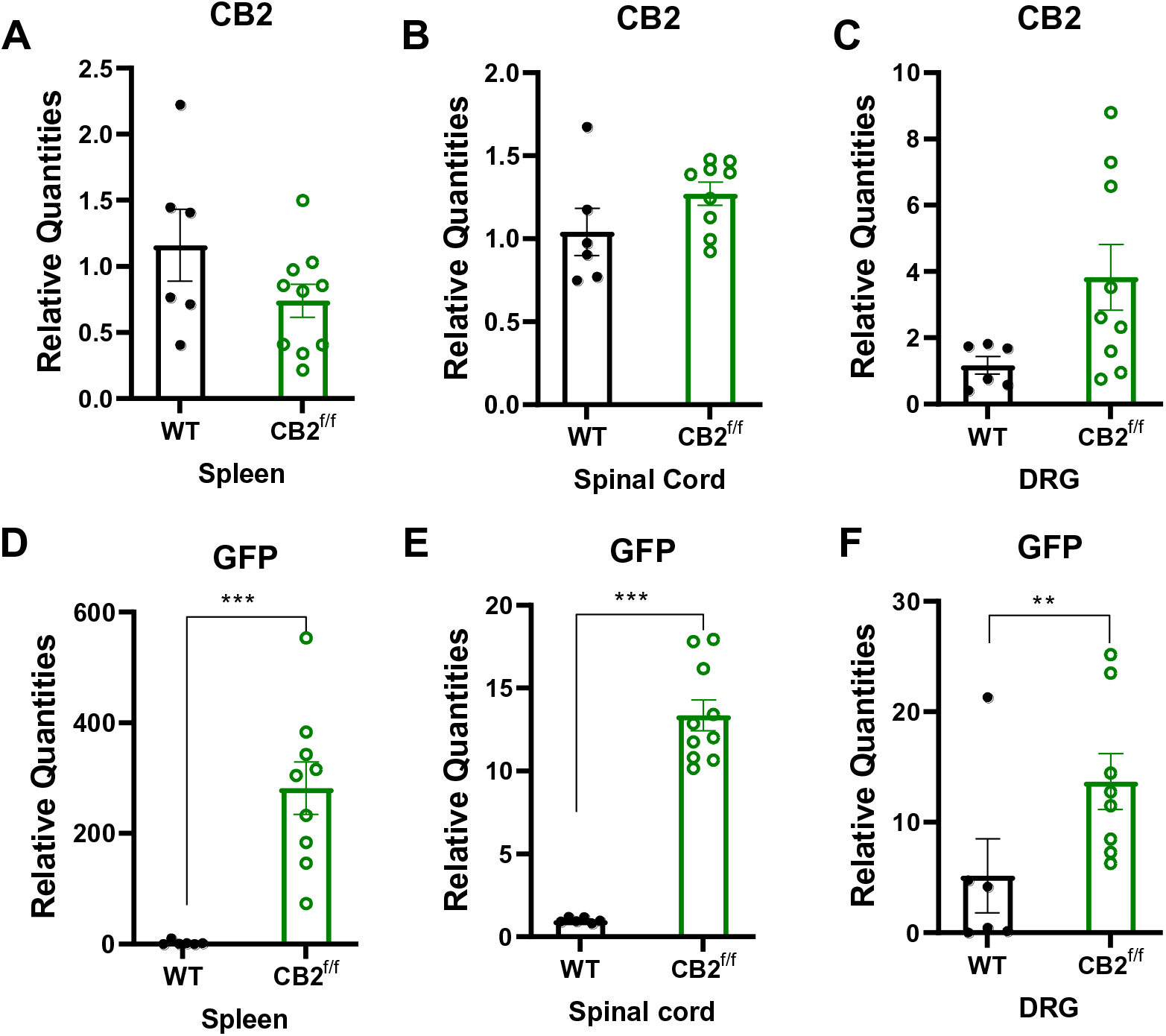
CB2 and GFP mRNA are detectable within nociceptive circuitry. CB2 and GFP mRNAs are detectable in tissues implicated in nociceptive processing. CB2 mRNA was present at comparable levels between CB2^f/f^ and wild type animals in the spleen, spinal cord and DRG (A-C). GFP mRNA was detectable in the spleen, spinal cord and DRG of CB2^f/f^ mice (D-F). (N=6 for wildtype group, N=11 for CB2^f/f^, animals used for this study were age matched between 90 days 150 days. Values were calculated using the 2(^-ΔΔ^Ct) method. GAPDH was used as reference gene. In WT tissues, CT values for GFP were typically undefined and assigned an arbitrary CT value of 37 for comparison with values in reporter mice. Data are expressed as mean ± S.E.M. **p < 0.01, ***p<0.001, unpaired sample two tailed t-test.

#### Comparison of CB2 and GFP mRNA expression levels in DRG, paw skin, lumbar spinal cord and spleen derived from advillin^Cre/+^;CB2^f/f^ and advillin^Cre/-^;CB2^f/f^ mice

To ascertain whether primary sensory neuron CB2 receptors mediated anti-allodynic effects of CB2 agonists, we generated advillin^Cre/+^;CB2^f/f^ conditional knockout mice. CB2 receptors were conditionally deleted from peripheral sensory neurons by crossing CB2^f/f^ mice with advillin^+/cre^ mice (15). In DRG, CB2 mRNA expression levels were lower in advillin^Cre/+^;CB2^f/f^ compared to advillin^Cre/-^;CB2^f/f^ mice (*p*=0.0014,Figure 9A). GFP mRNA was still detectable in DRG of advillin^Cre/+^;CB2^f/f^mice albeit at lower levels than that observed in advillin^Cre/-^;CB2^f/f^ mice (*p*=0.0056, Figure 9E). By contrast, CB2 mRNA expression levels did not differ between advillin^Cre/+^;CB2^f/f^ and advillin^Cre/-^;CB2^f/f^ mice in paw skin (Figure 9B), lumbar spinal cord (Figure 9C) or spleen (Figure 9D), indicating that insertion of GFP and deletion of CB2 gene in DRG did not alter the basal expression of CB_2_ in these tissues. As expected, GFP mRNA was detected in the paw skin, lumbar spinal cord and spleen of both advillin^Cre/-^;CB2^f/f^ and advillin^Cre/+^;CB2^f/f^ mice (Figure 9F-H). In addition, GFP mRNA was expressed at comparable level in paw skin, lumbar spinal cord and spleen of the advillin^Cre/-^;CB2^f/f^and advillin^Cre/+^;CB2^f/f^mice. Thus, advillin^Cre/+^;CB2^f/f^mice showed a selective deletion of CB2 and its EGFP reporter in DRG, but not in other tissues (spleen, spinal cord, paw skin), documenting the selectivity of our cKO mouse for deletion of primary sensory neuron CB2 expression levels in our study.

**Figure 9.**
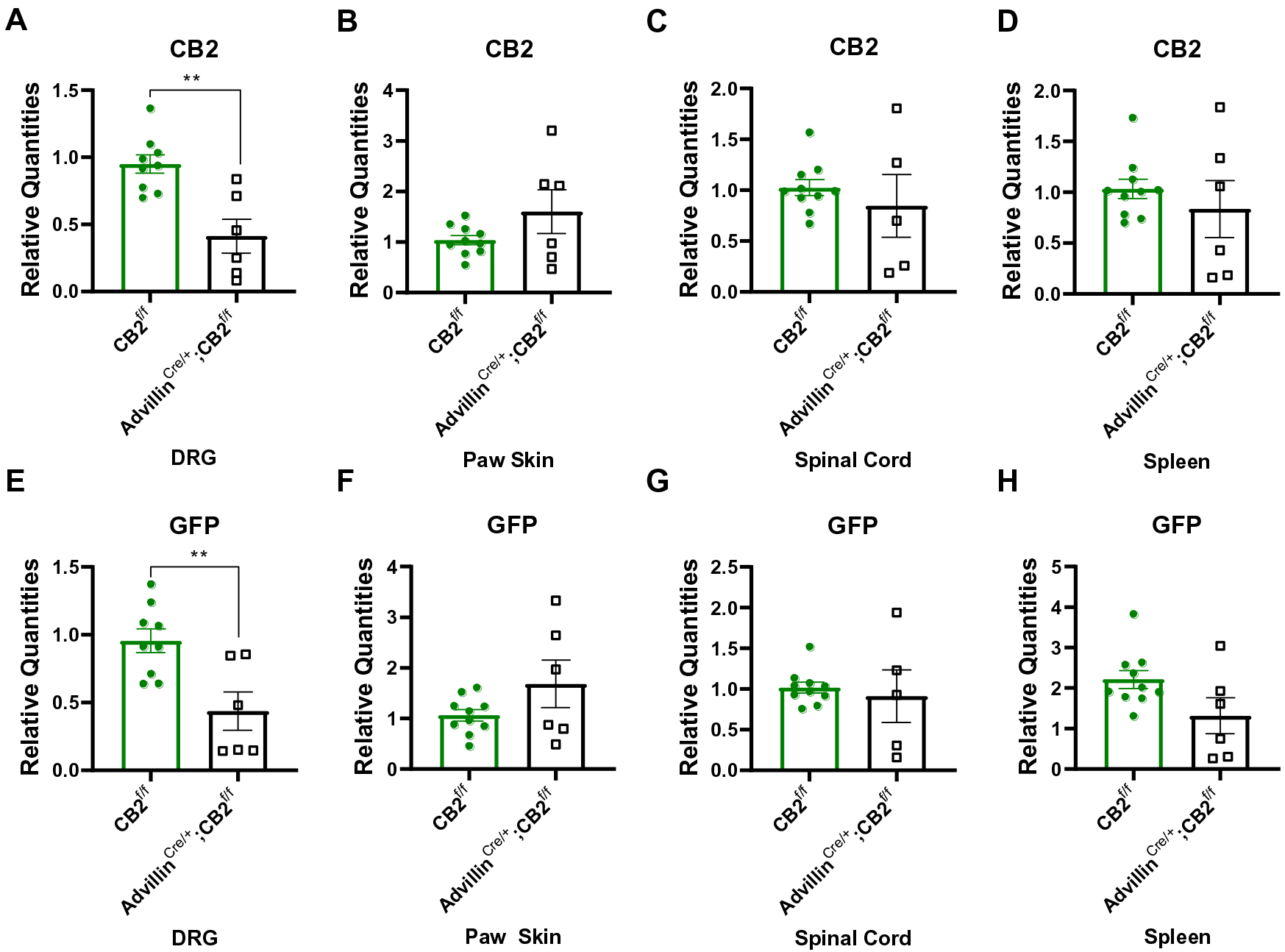
CB2 and GFP mRNA expression patterns in various tissues derived from advillin^Cre/+^;CB2^f/f^mice and CB2^f/f^ mice. Comparisons of (A-D) CB2 mRNA and (E-H) GFP mRNA in (A, B) DRG, (C, D) paw skin, (E, F) lumbar spinal cord and (G, H) spleen between advillin^Cre/+^;CB2^f/f^mice and CB2^f/f^ mice (n=6-10/group). Advillin^Cre/+^;CB2^f/f^mice display lower levels of CB2 and GFP mRNA in lumbar DRG relative to CB2^f/f^ mice, but no difference in expression levels between genotypes for either mRNA species in paw skin, spinal cord and spleen. Values were calculated using the 2(^-ΔΔ^Ct) method. GAPDH was used as reference gene. Data are expressed as mean ± S.E.M. **p < 0.01, unpaired sample two tailed t-test.

### CB2 agonists fail to alleviate ddC-evoked mechanical and cold allodynia in advillin^Cre/+^;CB2^f/f^ mice

Next, we assessed the impact of removing CB2 receptors from peripheral sensory neurons on the antinociceptive efficacy of LY2828360 and AM1710. Both AM1710 (Figure 10A,B) and LY2828360 (Figure 10C,D) alleviate ddC-evoked mechanical and cold hypersensitivity in male and female CB2^f/f^ mice. Specifically, AM1710 decreased ddC-evoked mechanical (Figure 10A) and cold (Figure 10B) hypersensitivity at 1 mg/kg (p<0.002 for all comparisons), 5 mg/kg (p<0.0001 for all comparisons), and 10 mg/kg (p<0.0001 for all comparisons), but not at 0.1 mg/kg (p>0.9 for all comparison). By contrast, AM1710 failed to alter mechanical or cold hypersensitivity in advillin^Cre/+^;CB2^f/f^mice at any dose (p>0.2 for all comparisons; Figure 10A,B). In the same subjects, morphine (10 mg/kg, i.p.) and gabapentin (50 mg/kg, i.p.) reversed ddC-induced mechanical and cold allodynia in all groups (p<0.0001 for all comparisons; Figure 10A,B). Similarly, LY2828360 decreased ddC-evoked mechanical (Figure 10C) and cold (Figure 10D) allodynia in CB2^f/f^ mice. In male and female CB2^f/f^ mice, LY2828360 attenuated ddC-evoked mechanical and cold hypersensitivity at 0.3 mg/kg (p<0.01 for all comparisons), 1 mg/kg (p<0.0001 for all comparisons) and 3 mg/kg (p<0.0001 for all comparisons), but not at 0.03 mg/kg (p>0.9 for all comparisons; Figure 10C). By contrast, LY2828360 did not alter ddC-evoked mechanical hypersensitivity in advillin^Cre/+^;CB2^f/f^ mice, at any dose (p>0.6 for all comparisons) (Figure 10C). Morphine (10 mg/kg, i.p.) and gabapentin (50 mg/kg, i.p.) reversed ddC-evoked mechanical hypersensitivity in both sexes and genotypes (p<0.0001 for all comparisons; Figure 10A-D). Vehicle treatment did not alter levels of mechanical or cold responsiveness in either sex or genotype (data not shown).

**Figure 10.**
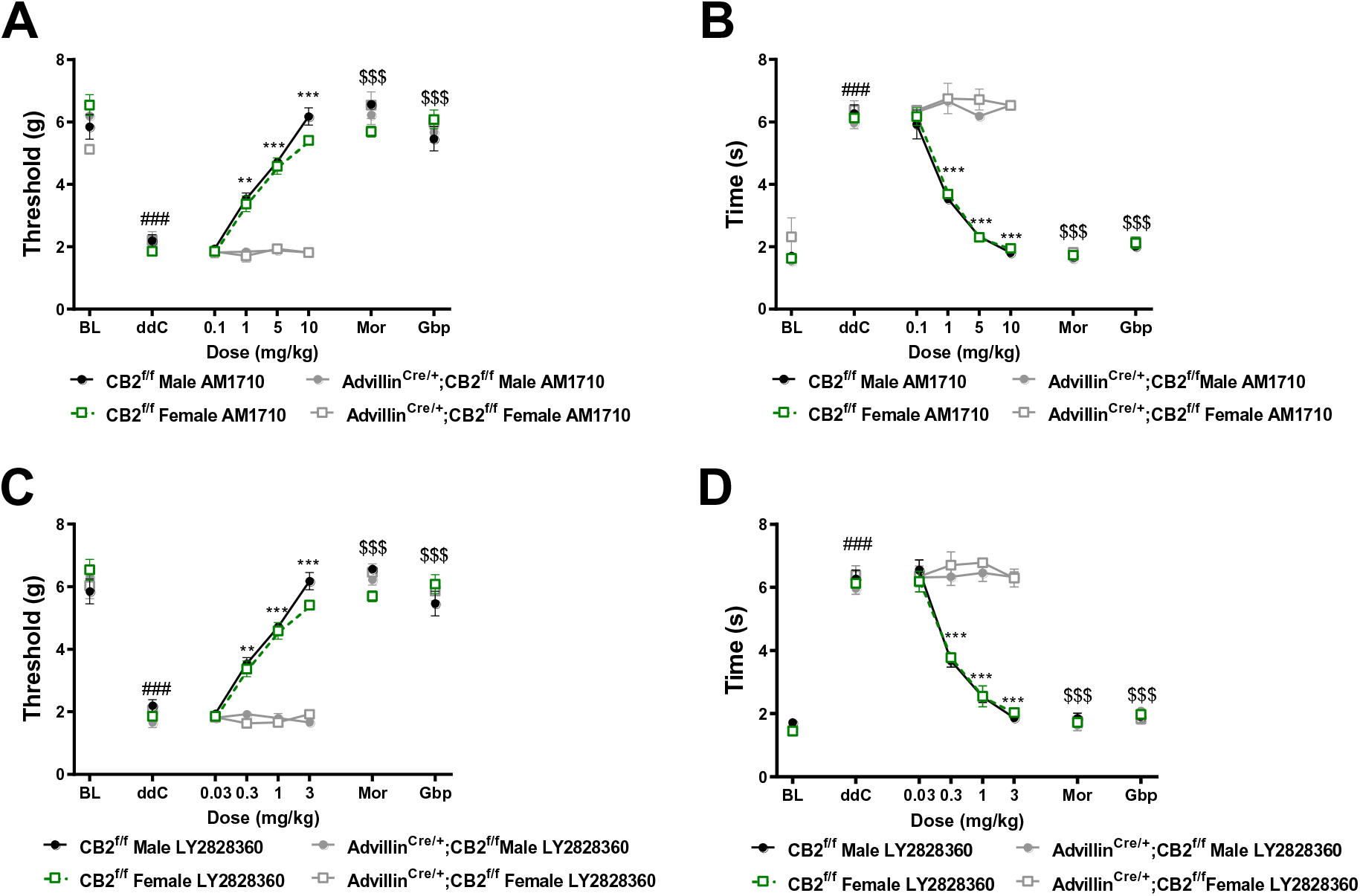
Removal of CB2 receptors from peripheral sensory neurons eliminates the antinociceptive efficacy of CB2 receptor agonists. AM1710 (A,B) and LY2828360 (C,D) dose dependently alleviate ddC-evoked mechanical and cold allodynia in CB2^f/f^ mice, but fail to alter ddC-induced behavioral hypersensitivities in advillin^Cre/+^;CB2^f/f^mice. The reference analgesics morphine and gabapentin reverse ddC-evoked mechanical (A) and cold (B) hypersensitivity in CB2^f/f^ and advillin^Cre/+^;CB2^f/f^ mice. Main effects of drug treatment for AM1710 (mechanical: F_7,112_ = 245.9, p<0.0001; cold: F_7,112_ = 384.7, p<0.0001), and LY2828360 (mechanical: F_7,112_ = 286.5, p<0.0001; cold: F_7,112_ = 414.9, p<0.0001), genotype for AM1710 (mechanical: F_3,16_ = 39.38, p<0.0001; cold: F_3,16_ = 46.6, p<0.0001) and LY2828360 (mechanical: F_3,16_ = 71.86, p<0.0001; cold: (F_3,16_ = 54.81, p<0.0001) and drug x genotype interactions for AM1710 (mechanical: F_21,112_ = 17.2, p<0.0001; cold: F_21,112_ = 37.66, p<0.0001) and LY2828360 (mechanical: F_21,112_ = 19.17, p<0.0001; cold: F_21,112_ =36.35, p<0.0001). ^###^p<0.0001 vs. pre-ddC baseline, **p<0.01, ***p<0.0001 male and female CB2^f/f^ vs. ddC, ^$$$^p<0.0001 all groups vs. ddC. Two-way ANOVA, Tukey multiple comparisons test post-hoc. (N= 5 per group). Mor: morphine, Gbp: gabapentin.

### LY2828360 prevents the development of tolerance to morphine’s antinociceptive efficacy in male CB2^f/f^ mice, and to a lesser extent in female CB2^f/f^ mice, whereas removal of CB2 receptors from peripheral sensory neurons eliminates these effects

In male mice, morphine (10 mg/kg i.p.) alleviated ddC-evoked mechanical (Figure 11A) and cold allodynia (Figure 11B) relative to vehicle treatment on days 1 (p<0.001) and 3 (p<0.01) of chronic dosing, but efficacy was absent on days 6-12 of repeated dosing (p>0.3 for all comparisons); these observations are consistent with development of morphine tolerance. LY2828360 co-administration (0.03 or 0.1 mg/kg, i.p.) with morphine (10 mg/kg, i.p.) prevented the development of tolerance to morphine’s antinociceptive efficacy (Figure 11A). Mechanical paw withdrawal thresholds were higher in mice treated with either dose of LY2828360 in combination with morphine on day 1-12 (p<0.0001) relative to mice receiving vehicle and on days 3-12 of administration relative to mice receiving morphine alone (p<0.001 for each comparison). Cold responsiveness varied slightly between the two doses of LY2828360 (Figure 11B). Mice treated with the high dose of LY2828360 (0.1 mg/kg) in conjunction with morphine displayed lower levels of cold responsiveness (p<0.01) on day 1 of administration relative to mice treated with the low dose of LY2828360 (0.03 mg/kg). Both doses of LY2828360 suppressed the development of morphine tolerance, producing sustained relief of ddC-evoked cold hypersensitivity on days 3 (p<0.01) and 6-12 of administration (p<0.001) relative to mice treated with either vehicle or morphine alone (Figure 11B).

**Figure 11.**
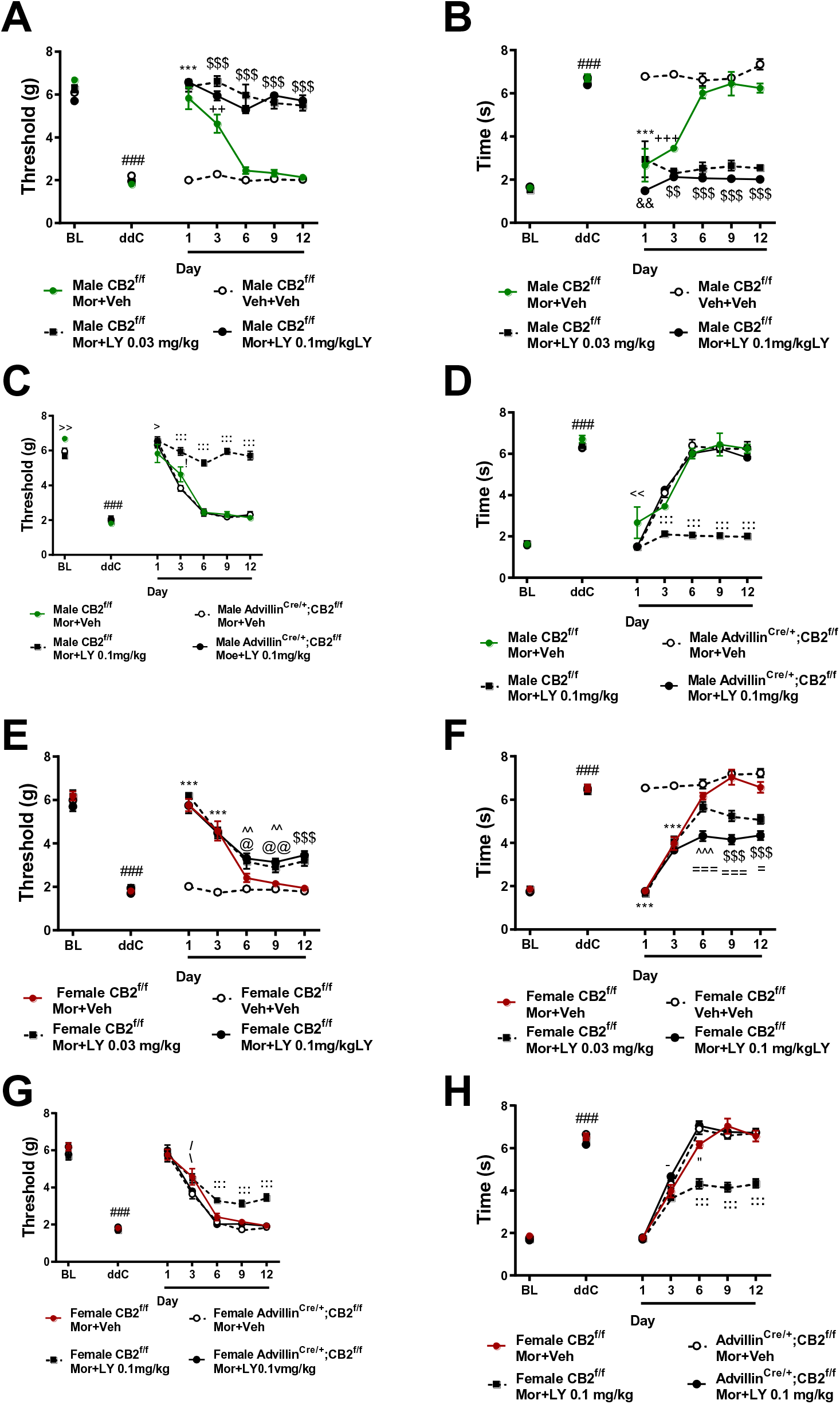
Co-administration of LY2828360 with morphine prevents the development of tolerance to morphine’s antinociceptive effects in male, and to a lesser extent in female CB2^f/f^ mice but not in advillin^Cre/+^;CB2^f/f^ mice. Morphine (10 mg/kg i.p.) alleviated ddC-evoked mechanical (A, C, E, G) and cold (B, D, F, H) responsiveness on days 1 and 3 of administration. Co-administration of LY2828360 (0.03 mg/kg or 0.1 mg/kg i.p.) with morphine (10 mg/kg i.p.) prevented the development of tolerance, producing sustained suppressions of ddC-evoked mechanical (A,C) or cold (B,D) allodynia on days 1-12 of administration in male CB2^f/f^ mice but not male advillin^Cre/+^;CB2^f/f^ mice. Co-administration of LY2828360 (0.03 mg/kg or 0.1 mg/kg i.p.) with morphine (10 mg/kg i.p.) lessened the development of tolerance in CB2^f/f^ mice, though to a lesser extant than that observed in male CB2^f/f^ mice (E-H). The tolerance sparing effects of LY2828360 were absent in female; CB2^f/f^ mice (G,H).Male CB2^f/f^ mice co-administration schedule: main effects of ddC (mechanical: F_6,192_ = 162.2 p<0.0001; cold: F_6,192_ = 131.8, p<0.0001), LY2828360 treatment (mechanical: F_3,32_ =159.2, p<0.0001; cold: F_3,32_ = 174.2, p<0.0001) and ddC x LY2828360 interaction (mechanical: F_18,192_ = 32.51, p<0.0001; cold: F_18,192_ = 24.9, p<0.0001) were observed. Male CB2^f/f^ vs. advillin^Cre/+^;CB2^f/f^ co-administration schedule main effects of drug treatment (mechanical: F_6,126_ = 243.3, p<0.0001; cold: F_6,162_ = 212.9, p<0.0001), genotype (mechanical: F_3,27_ = 78.5, p<0.0001; cold: F_3,27_ = 139.5, p<0.0001) and treatment x genotype interactions (mechanical: F_18,162_ = 22.11, p<0.0001; cold: F_18,162_ = 20.42, p<0.0001) were observed. Male mice reversal schedule: main effects of ddC (mechanical: F_6,180_ = 201, p<0.0001; cold: F_6,180_ = 147, p<0.0001), LY2828360 treatment (mechanical: F_3,30_ = 90.69, p<0.0001, cold: F_3,30_ = 124.5, p<0.0001) and ddC x LY2828360 interaction (mechanical: F_18,180_ = 32.5, p<0.0001, cold: F_18,180_ = 23.47, p<0.0001) were observed. Female CB2^f/f^ mice co-administration schedule: main effects of ddC (mechanical: F_6,168_ = 210.6, p<0.0001; cold: F_1,168_ = 403.3, p<0.0001), LY2828360 treatment (mechanical: F_3,28_ = 52.67, p<0.0001; cold: F_3,28_ = 111.4, p<0.0001) and ddC x LY2828360 interaction (mechanical: F_18,168_ = 14.68, p<0.0001; cold: F_18,168_ = 31.51, p<0.0001) were observed. Female CB2^f/f^ vs. advillin^Cre/+^;CB2^f/f^ co-administration dosing schedule main effects of drug treatment (mechanical: F_6,156_ = 322.4, p<0.0001; cold: F_6,156_ = 618.3, p<0.0001), genotype (mechanical: F_3,26_ = 9.351, p<0.001; cold: F_3,26_ = 37.18, p<0.0001) and treatment x genotype interactions (mechanical: F_18,156_ = 4.47, p<0.0001; cold: F_18,156_ = 15.85, p<0.0001) were observed. ^###^p<0.001 vs. baseline, ***p<0.001 all groups vs. veh+veh, ^$$$^p<0.001, ^$$^p<0.01 Mor+LY 0.1mg/kg and Mor+ LY 0.03 mg/kg vs. all other groups, ^+++^p<0.001, ^++^p<0.01 MOR+Veh vs. Veh+Veh, ^&&^p<0.01, ^&^p<0.05 Mor+LY0.1mg/kg vs. Mor+LY 0.03 mg/kg, ^@@^p<0.01, ^@^p<0.05 Mor+LY 0.1mg/kg vs. Mor+Veh, ^^^^p<0.01 Mor+LY 0.1 mg/kg and Mor+LY 0.03 mg/kg vs. Veh+Veh, ^===^p<0.001, ^=^p<0.05 Mor+LY 0.1 mg/kg vs. all other groups, ^%^p<0.05 Mor+LY 0.03 mg/kg vs. Veh, ^--^p<0.01 Mor+LY 0.03 mg/kg vs. Veh+Veh and Mor+LY 0.1 mg/kg. ^:::^p<0.001, ^::^p<0.01, :p<0.05 CB2^f/f^ Mor+LY vs. all other groups, ^>^p<0.05 CB2^f/f^ Mor+LY vs. CB2^f/f^ Mor+Veh, ^!^p<0.05 CB2^f/f^ Mor+Veh vs. Adv cre+;CB2 f/f Mor+LY, ^<<^p<0.01, ^<^p<0.05 CB2^f/f^ MOR+Veh vs. all other groups, ^\^p<0.05 CB2^f/f^ MOR+LY vs. vs. advillin^Cre/+^;CB2^f/f^ Mor+VEH, ^-^ p<0.05 CB2^f/f^ vs. advillin^Cre/+^;CB2^f/f^ Mor+LY, ^“^p<0.05 CB2^f/f^ vs. **^+^**;CB2^f/f^, ^∼^p<0.05 CB2^f/f^ Mor+LY vs. advillin^Cre/+^;CB2^f/f^ Mor+LY, ^/^p<0.05 CB2^f/f^ Mor+Veh vs. advillin^Cre/+^;CB2^f/f^ Mor+LY.Two-way ANOVA, Tukey Multiple comparisons test post-hoc. (N=6-10 per group). Black bars indicate period of LY and morphine co-administration. Mor: morphine, LY: LY2828360, Veh: vehicle.

In female mice, morphine alleviated ddC-evoked mechanical and cold allodynia on days 1 and 3 of administration (p<0.001 for all comparisons), but not on days 6-12 of administration (p>0.05 for all comparisons) (Figure 11E, F). LY2828360 (0.03 mg/kg or 0.1 mg/kg i.p.) co-administration prolonged the antinociceptive efficacy of morphine. Female mice treated with either dose of LY2828360 displayed higher mechanical withdrawal thresholds on day 6 and 9 of co-administration (p<0.01) relative to female mice receiving vehicle, and relative to all groups on days and 12 (p<0.0001 for each comparison) (Figure 11E). The high dose of LY2828360 in conjunction with morphine also elevated paw withdrawal thresholds on days 6-12 (p<0.05 for all comparisons) relative to female mice receiving morphine alone. Similarly, both doses of LY2828360 reduced cold responsiveness on day 6 relative to vehicle (p<0.001 for each comparison) and on days 9 and 12 relative to all other groups (p<0.001 for each comparison, Figure 11F). LY2828360 dose-dependently lessened the development of tolerance to morphine’s ability to suppress cold hypersensitivity. Female mice treated with the high dose of LY2828360 in conjunction with morphine displayed lower levels of cold responsiveness on days 6-12 of administration relative to female mice receiving the low dose of LY2828360 in conjunction with morphine (p<0.05 for all comparisons, Figure 11F).

In both genotypes, ddC induced hypersensitivity to mechanical and cold stimulation (p<0.0001 for all comparisons) (Figure 11C, D, G, H). Morphine (10mg/kg, i.p.) co-administered with LY2828360 (0.1 mg/kg, i.p.) increased mechanical paw withdrawal thresholds (Figure 11C) and lowered cold responsiveness (Figure 11D) on days 3-12 (p<0.001 for all comparisons) in male (Figure 11C, D) and female (Figure 11G, H) CB2^f/f^ mice, but not in advillin^Cre/+^;CB2^f/f^ mice (Figure 11C, D, G, H). Thus, removal of CB2 receptors from peripheral sensory neurons eliminated the efficacy of CB2 agonists in sparing morphine tolerance. Transient differences in mechanical and cold allodynia were observed between groups throughout the testing period. Male CB2^f/f^ mice treated with morphine alone displayed modestly higher mechanical withdrawal thresholds at baseline (p=0.003), and lower mechanical withdrawal thresholds following acute drug treatment (dose 1; p=0.04) relative to CB2^f/f^ mice treated with morphine in combination with LY2828360. Male CB2^f/f^ mice treated with morphine alone also displayed higher mechanical paw withdrawal thresholds following dose 3 (p=0.04) relative to male advillin^Cre/+^;CB2^f/f^ mice treated with morphine in combination with LY2828360. Male CB2^f/f^ mice treated with morphine displayed higher levels of cold responsiveness following dose 1 relative to all other groups (p<0.01 for all comparisons).

In female CB2^f/f^ mice, morphine (10 mg/kg, i.p.) in combination with LY2828360 (0.1 mg/kg, i.p.) increased mechanical withdrawal thresholds relative to female advillin^Cre/+^;CB2^f/f^ mice receiving morphine alone on day 3 (p=0.02), and relative to all other groups on days 6-12 (p<0.01 for all comparisons). Female CB2^f/f^ mice treated with morphine alone displayed higher mechanical paw withdrawal thresholds on day 3 of administration relative to female advillin^Cre/+^;CB2^f/f^ mice treated with morphine alone or in combination with LY2828360 (p<0.05 for each comparison; Figure 11G).

### LY2828369 reverses established morphine tolerance in male, and to a lesser extent in female CB2^f/f^ mice, whereas removal of CB2 receptors from peripheral sensory neurons eliminates the tolerance reversing effects LY2828360

In male CB2^f/f^ mice receiving the reversal dosing schedule, prior to initiation of LY2828360 dosing (0.1 mg/kg i.p., days 7-12 of administration), morphine (10 mg/kg i.p.) produced initial antinociceptive efficacy, alleviating ddC-evoked mechanical (Figure 12A) and cold (Figure 12B) allodynia on days 1 and 3 of administration (p<0.0001 for each comparison). By day 6 of administration, morphine’s antinociceptive efficacy was no longer observed (p>0.3 for all comparisons), consistent with development of tolerance. Treatment with LY2828360 (0.03 or 0.1 mg/kg i.p.) beginning on day 7 of morphine administration reversed established morphine tolerance, alleviating ddC-evoked mechanical (Figure 12A) and cold (Figure 12B) hypersensitivity on days 9 and 12 of administration relative to male mice treated with morphine alone or vehicle (p<0.0001 for all comparisons). Male mice treated with LY2828360 (0.1 mg/kg i.p. on days 7-12 of administration), however, displayed modestly lower levels of mechanical hypersensitivity on days 6 (p<0.05) and 9 (p<0.01) (Figure 12A) and lower levels of cold allodynia on day 9 (p<0.05) of administration (Figure 12B) relative to male mice receiving the low dose of LY2828360. Neither dose of LY2828360 alone altered mechanical or cold responsiveness in the absence of morphine (data not shown).

**Figure 12.**
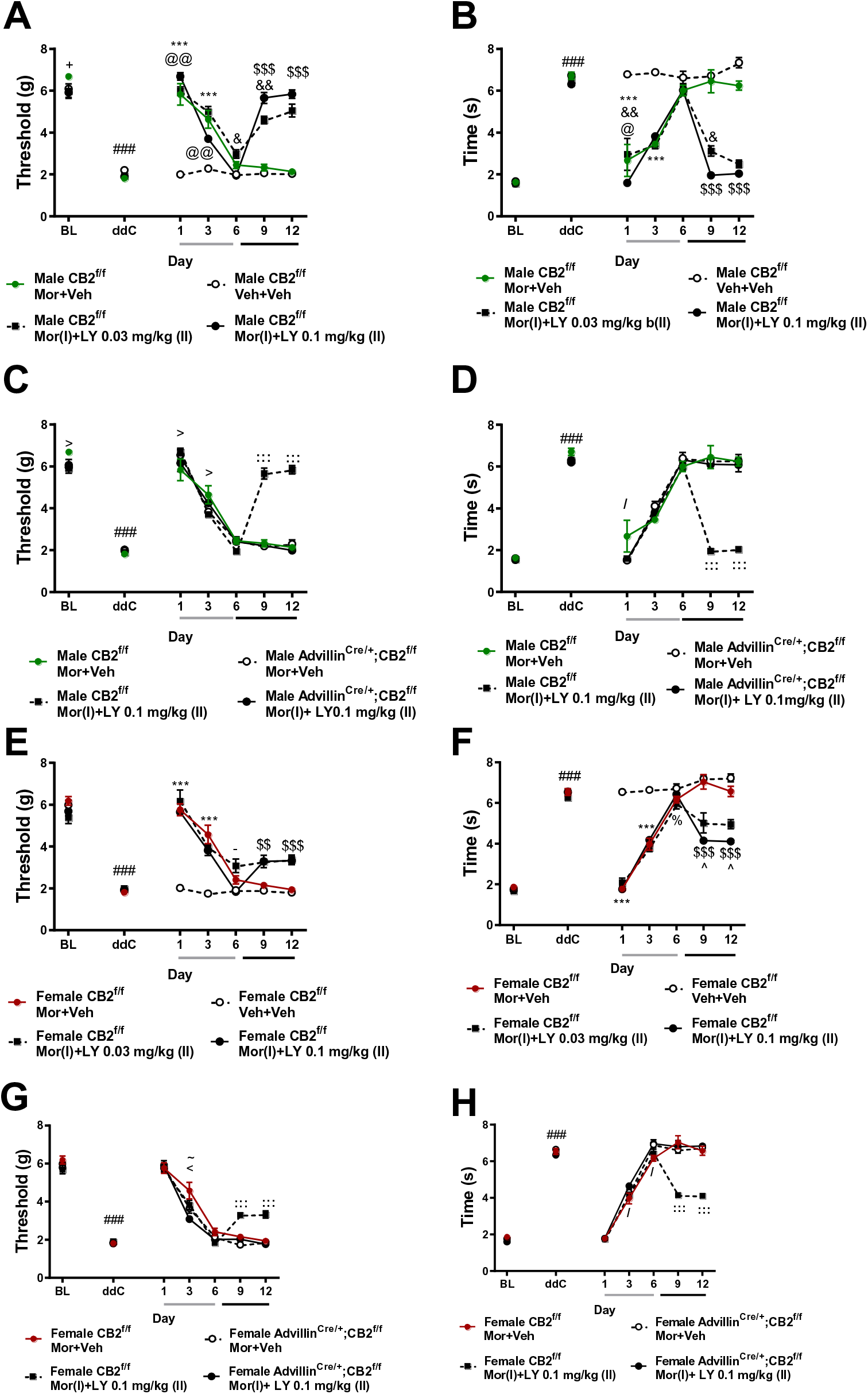
LY2828360 reverses established tolerance to the anti-allodynic effects of morphine in male, and to a lesser extent in female CB2^f/f^ mice, but not in advillin^Cre/+^;CB2^f/f^ mice. In all groups receiving morphine (10 mg/kg i.p.) alone on days 1-6 of administration, tolerance rapidly develops to morphine’s antinociceptive efficacy (A-H). In CB2^f/f^ mice, following initiation of LY2828360 dosing (0.03 mg/kg or 0.1 mg/kg i.p.) in conjunction with morphine on days 7-12, the antinociceptive efficacy of morphine returned, alleviating ddC-evoked mechanical (A, C, E, G) and cold (B, D, F, H) hypersensitivity on days 9-12 of administration. LY2828360 produced a reversal of established morphine tolerance in male, and to a lesser extent in female CB2^f/f^ mice (A-H), but not in advillin^Cre/+^;CB2^f/f^ mice of either sex (C,D,G,H). Male CB2^f/f^ mice reversal schedule: main effects of ddC (mechanical: F_6,180_ = 201, p<0.0001; cold: F_6,180_ = 147, p<0.0001), LY2828360 treatment (mechanical: F_3,30_ = 90.69, p<0.0001, cold: F_3,30_ = 124.5, p<0.0001) and ddC x LY2828360 interaction (mechanical: F_18,180_ = 32.5, p<0.0001, cold: F_18,180_ = 23.47, p<0.0001) were observed. Male CB2^f/f^ vs. advillin^Cre/+^;CB2^f/f^ reversal dosing schedule main effects of drug treatment (mechanical: F_6,156_ = 243.6, p<0.0001; cold: F_6,156_ = 221.7, p<0.0001), genotype (mechanical: F_3,26_ = 20.89, p<0.0001; cold: F_3,26_ = 43.37, p<0.0001) and treatment x genotype interactions (mechanical: F_18,156_ = 20.93, p<0.0001; cold: F_18,156_ = 18.11, p<0.0001) were observed. Female CB2^f/f^ mice reversal schedule: main effects of ddC (mechanical: F_6,162_ = 178.5, p<0.0001; cold F_6,162_ = 362.7, p<0.0001), LY2828360 treatment (mechanical: F_3,27_ = 60.03, p<0.0001; cold: F_3,27_ = 96.56, p<0.0001), and ddC x LY2828360 interactions (mechanical: F_18,162_ = 15.06, p<0.0001; cold: F_18,162_ = 29.34, p<0.0001) were observed. Female CB2^f/f^ vs. advillin^Cre/+^;CB2^f/f^ reversal dosing schedule main effects of drug treatment (mechanical: F_6,156_ = 377.4,p<0.0001; cold: F_6,156_ = 617.3, p<0.0001), genotype (mechanical: F_3,26_ = 6.8, p<0.001, cold:F_3,26_ = 26.18, p<0.0001) and treatment x genotype interaction (mechanical: F_18,156_ = 6.281, p<0.0001; cold: F_18,156_ = 14.87, p<0.0001) were observed. ^###^p<0.001 vs. baseline, ^:::^p<0.001, ^::^p<0.01, :p<0.05 CB2^f/f^ Mor+LY vs. all other groups, ^>^p<0.05 CB2^f/f^ Mor+LY vs. CB2^f/f^ Mor+Veh, ^!^p<0.05 CB2^f/f^ Mor+Veh vs. Adv^Cre/+;^CB2 f/f Mor+LY, ^<<^p<0.01, ^<^p<0.05 CB2^f/f^ Mor+Veh vs. all other groups, ^\^p<0.05 CB2^f/f^ Mor+LY vs. vs. advillin^Cre/+^;CB2^f/f^ Mor+VEH, ^-^p<0.05 CB2^f/f^ vs. advillin^Cre/+^;CB2^f/f^ Mor+LY, ^“^p<0.05 CB2^f/f^ vs. advillin^Cre/+^;CB2^f/f^, ^∼^p<0.05 CB2^f/f^ Mor+LY vs. advillin^Cre/+^ ;CB2^f/f^ Mor+LY, ^/^p<0.05 CB2^f/f^ Mor+Veh vs. advillin^Cre/+^;CB2^f/f^ Mor+LY.Two-way ANOVA, Tukey Multiple comparisons test post-hoc. (N=6-10 per group). Black bars indicate period of LY and morphine co-administration. For reversal condition, grey bar indicates period of vehicle+morphine administration (days 1-6), black bar indicates period of LY+morphine co-administration (days 7-12). Mor: morphine, LY: LY2828360, Adv: advillin, Veh: vehicle.

In female mice receiving the dosing schedule used to reverse established tolerance, morphine alleviated ddC-evoked mechanical and cold allodynia on days 1 and 3 (p<0.0001 for all comparisons) in all morphine-treated groups (Figure 12E). By day 6, the antinociceptive effects of morphine were largely absent, consistent with the development of tolerance. Initiation of LY2828360 treatment on day 7 of morphine administration reversed established tolerance to morphine’s antinociceptive efficacy. Both doses of LY2828360 (0.03 mg/kg or 0.1 mg/kg i.p. on day 7-12) in conjunction with morphine increased mechanical paw withdrawal thresholds (Figure 12E), and lowered levels of cold responsiveness (Figure 12F) on days 9 and 12 relative to all other groups (p<0.001 for all comparisons). Additionally, the high dose of LY2828360 was more efficacious than the low dose in female mice, producing reduced cold responsiveness on days 9 and 12 of administration relative to all other groups (p<0.05 for each comparison; Figure 12F). In female mice, LY2828360 did not reliably alter ddC-evoked allodynia in the absence of morphine (data not shown).

In male and female CB2^f/f^ mice receiving LY2828360 (0.1 mg/kg, i.p.) co-administered with morphine on days 7-12, mechanical withdrawal thresholds were higher and cold responsiveness was lower on days 9 and 12 (p<0.001 for each comparison) relative to CB2^f/f^ mice receiving morphine alone or advillin^Cre/+^;CB2^f/f^ mice receiving either treatment (Figure 12C-D, G-H). In male CB2^f/f^ mice, prior to initiation of reversal dosing with LY2828360 all groups displayed similar levels of mechanical and cold allodynia with a few exceptions. Male CB2^f/f^ mice treated with morphine alone displayed higher mechanical withdrawal thresholds (Figure 12A, C) at baseline (p=0.04) and after dose 3 (p=0.01) relative to male CB2^f/f^ mice treated with the reversal dosing schedule of LY2828360, and lower mechanical withdrawal thresholds following the first injection (p=0.02) relative to male CB2^f/f^ mice treated with the reversal dosing schedule of LY2828360. Male CB2^f/f^ mice treated with morphine alone also displayed higher levels of cold responsiveness following dose 1 relative to all other groups (p<0.05 for all comparisons; Figure 12B, D). Female CB2^f/f^ and advillin^Cre/+^;CB2^f/f^ mice treated with morphine alone or with the reversal dosing schedule displayed roughly equivalent mechanical (Figure 12G) and cold (Figure 12H) responses prior to the initiation of reversal dosing with LY2828360 on day 7. On day 3 of administration, female CB2^f/f^ mice treated with morphine alone displayed higher mechanical withdrawal thresholds relative to all other groups (p<0.05 for each comparison). Prior to initiating the reversal dosing regimen of LY2828360 and morphine, morphine increased mechanical withdrawal thresholds (Figure 12G) and reduced cold responsiveness (Figure 12H) in female CB2^f/f^ mice relative to female advillin^Cre/+^;CB2^f/f^ mice in the same treatment group on day 3 (p=0.03). On day 6, female CB2^f/f^ mice treated with morphine alone displayed lower levels of cold responsiveness relative to both groups of female advillin^Cre/+^;CB2^f/f^ mice (p<0.05 for all comparisons). In male and female CB2^f/f^ and advillin^Cre/+^;CB2^f/f^ mice treated with vehicle in lieu of morphine, LY2828360 did not alter mechanical or cold responsiveness (p>0.1 for all comparisons, data not shown). These results indicate that, in male and female mice, removal of CB2 from peripheral sensory neurons eliminates the ability of LY2828360 to reverse established morphine tolerance.

## Discussion

The present studies provide the first evidence that CB2 receptors on peripheral sensory neurons are necessary for the antinociceptive effects of CB2 agonists. They also support a previously unrecognized role for primary sensory CB2 receptors in mediating the ability of CB2 agonists to both spare and reverse opioid antinociceptive tolerance. These studies further validate CB2 receptors as a viable therapeutic target for the management of ATN-associated pain. In our studies, two structurally distinct CB2 agonists, AM1710 and LY2828360, alleviated ddC-evoked mechanical and cold hypersensitivity at a wide range of doses, without altering mechanical or cold responsiveness in the absence of ddC. The antinociceptive efficacy of AM1710 and LY2828360 was absent in mice globally lacking CB2, indicating the pain-relieving effects of both compounds are mediated by CB2 receptors. In these same CB2 KO mice, morphine completely reversed ddC-evoked mechanical and cold hypersensitivity, further indicating that the lack of antinociceptive efficacy of LY2828360 and AM1710 is not due to some gross alteration of nociceptive circuitry resulting from CB2 deletion. The present studies add to the previous body of literature from preclinical to clinical research suggesting that the cannabinoid system may be targeted to efficaciously alleviate HIV-associated sensory neuropathy. Cannabis consumption in HIV-infected patient populations provides a range of beneficial effects, from alleviating HIV-associated pain (3–5, 17), to increasing appetite and reducing nausea and psychiatric symptoms such as anxiety and depression (5). Evidence from preclinical research also supports the therapeutic potential of targeting the cannabinoid system for alleviating both primary sources implicated in the development of HIV-associated pain. Mixed CB1/CB2 agonists (18), and selective CB2 agonists have demonstrated efficacy in alleviating pain produced by administration of the HIV-associated toxic viral protein gp120 (19). Mixed CB1/CB2 agonists (20), exogenously administered endocannabinoids (21), and the *Cannabis sativa* terpene β-caryophyllene which is thought to exert its pharmacological effects via CB2 receptors, similarly have proven efficacious in alleviated ATN-associated pain (9).

CB2 receptor stimulation alleviates allodynia in preclinical pain models without producing the problematic side effects associated with CB1 receptor stimulation (7), including the development of tolerance (8, 22, 23), cannabimimetic side effects or physical dependence (8). In the current study, AM1710 and LY2828360 produced sustained efficacy in reducing ddC-evoked mechanical and cold hypersensitivity following 12 days of chronic dosing, at similar levels between male and female mice, whereas mice rapidly developed tolerance to the antinociceptive effects of morphine. The results of the current study support the possibility that targeting CB2 represents a superior therapeutic target. In addition to tolerance developing to the antinociceptive effects and the high risk for abuse and addiction associated with opioids such as morphine, opioids may exacerbate HIV-induced neuroinflammation and infectivity (24). MOR activation on glial cells potentiates the release of proinflammatory cytokines and chemokines induced by HIV protein byproducts such as Tat (25–27), indicating MOR agonist administration to HIV-infected individuals has the potential to worsen HIV-induced neuroinflammation and propagate inflammation through the recruitment of additional immune cells to sites of inflammation. Furthermore, morphine administration also upregulates the co-receptors HIV uses to enter and infect immune cells, such as CCR5 (28–30) and CXCR4 (31), which is implicated in the development of HIV-associated neuropathic pain (32, 33). MOR agonist administration in HIV-infected patient populations therefore has the potential to worsen HIV-induced neuroinflammation, HIV-infectivity and HIV-associated neuropathic pain. In contrast, CB2 stimulation may be protective against HIV-induced neurotoxicity/neuroinflammation (34, 35).

The present studies provide the first evidence that CB2 expression on peripheral sensory neurons may be responsible for the antinociceptive efficacy of CB2 agonists. Selective deletion of CB2 from excitatory neurons or peripheral sensory neurons completely eliminated the antinociceptive efficacy of two structurally distinct CB2 receptor agonists. Moreover, these effects were observed in both male and female mice. On the other hand, in advillin^Cre/+^;CB2^f/f^mice, two mechanistically distinct reference analgesics (morphine and gabapentin) still effectively eliminated ddC-induced mechanical and cold hypersensitivity. Additionally, the two CB2 receptor agonists used in the current studies exhibit differential levels of CNS availability. LY2828360 readily crosses the blood brain barrier, whereas AM1710 exhibits limited CNS penetration (36, 37). Given that both compounds displayed a similar ability to decrease ddC-induced mechanical and cold hypersensitivity suggests CB2 stimulation exerting antinociceptive effects via a peripheral site of action. There exists considerable controversy regarding the distribution of CB2 receptors within the central nervous system (CNS) (12). A growing number of reports indicate CB2 receptors may be present within the CNS, though the types of cells, and under what circumstances they may be expressed are still debated. Multiple examples have demonstrated the possibility of CB2 expression within the CNS (13, 38–54), with functions ranging from modulating emesis (38), hippocampal plasticity (41, 52), learning and memory (54), the rewarding effects of drugs of abuse (42, 44, 47) or the excitability of ventral tegmental dopaminergic neurons in general (42, 46), the descending control of pain (43), Parkinson’s disease (45), schizophrenia (48), anxiety (49), depression (50) and Alzheimer’s (13) disease. CB2 expression has also been demonstrated to increase in primary afferent neurons under pathological conditions (55–57). In the present studies, native GFP fluorescence and anti-GFP immunoreactivity were below the threshold for detection throughout the brain and sensory neurocircuitry including in the cell bodies of primary afferent neurons as well as the dorsal horn of the lumbar spinal cord, where primary afferent neurons terminate in the central nervous system. However, CB2 mRNA was detected in both lumbar DRG and lumbar spinal cord, albeit at low levels, suggesting immunohistochemical approaches may have insufficient sensitivity to detect protein expressed at such low levels. Both CB2 and GFP mRNA were decreased in the DRG of advillin^Cre/+^;CB2^f/f^ mice, but not in the spleen or spinal cord relative to CB2^f/f^ mice, indicating that deletion of CB2 receptors by advillin^Cre/+^ selectively decreased mRNAs for both markers of CB2 receptor transcription in the cell bodies of peripheral sensory neurons. Additionally, local administration of CB2 receptor agonists in the paw decrease mechanical and cold allodynia induced by the chemotherapeutic drug paclitaxel, and CB2^EGFP^ is detected in keratinocytes, Langerhans cells and dendritic cells within the epidermis (14). In fact, using the same CB2^f/f^ mouse used here, we recently showed that CB2^EGFP^ in Langerhans cells and keratinocytes are dynamically regulated by paclitaxel in peripheral paw tissue under conditions in which only scattered axonal labeling of large diameter fibers were observed (14). These latter effects are consistent with likely localization of CB2 on Aβ fibers whereas thinly myelinated and unmyelinated epidermal nerve fibers lacked CB2^EGFP^ labeling (14). CB2 agonist administered locally in the paw also suppressed paclitaxel-induced allodynia and increases spinal IL-10 mRNA levels (14). These observations raise the possibility that CB2 on the peripheral terminals of primary afferents may contribute to the antinociceptive effects of CB2 receptor agonists in ddC-induced neuropathic pain. Further studies are needed to determine whether GFP/CB2 can be detected in other areas on primary afferent neurons such as peripheral terminals in the paw skin, or whether more sensitive techniques such as RNAscope may provide insight into the tissues and cells expressing GFP and/or CB2 mRNA.

Recently, it was reported that global deletion of CB2 receptors or selective removal of CB2 from myeloid cells exacerbated the development of neuropathic pain induced by partial ligation of the sciatic nerve (58), whereas removal of CB2 from neuronal populations had no effect on the development of pain. However, this study did not examine the efficacy of CB2 agonists in ameliorating established pain in neuronal CB2-knockout mice. Furthermore, given previous reports indicating CB2 protein expression is induced following peripheral nerve injury (55, 57), these results do not preclude the possibility that CB2 in neuronal populations might contribute to antinociception after neuropathic pain has been established. In a subsequent report, removal of CB2 from neurons in mice expressing Cre recombinase under the *Synapsin I* promoter (Syn-Cre+:Cnr2^fl/fl^), but not Nav1.8+ primary afferents (Nav1.8-Cre+:-Cnr2^fl/fl^), modestly attenuated the ability of the CB2 receptor agonist JWH133 to alleviate mechanical hypersensitivity in a partial sciatic nerve injury model (59). This slight decrease in the antinociceptive efficacy of JWH133 was accompanied by an increase in JWH133 self-administration, suggesting an increase in spontaneous pain and indicating that despite increased intake of a CB2 agonist the ability of JWH133 to alleviate established neuropathic pain was blunted (but not eliminated) by removal of CB2 receptors from neurons. The sustained efficacy of JWH133 following removal of CB2 from Nav1.8+ nociceptors is interesting given the robust effects observed in the present studies. Discrepancies may be explained by the fact that, in the present study, CB2 was removed from *all* peripheral sensory neurons, not just Nav1.8+ nociceptive primary afferents. Additionally, the present study utilized a model of toxic neuropathy whereas a surgically-induced peripheral nerve injury model was used in the previous report. The presence of CB2 on peripheral sensory nerves indicated by the present study is further supported by previous reports demonstrating that *in vitro* CB2 receptor agonists produce an inhibitory effect on the activity of guinea pig and/ or human vagus nerve preparations to stimulation via prostaglandin E2 (guinea pig), hypertonic saline (guinea pig) or capsaicin (both) (60). Additionally, in cultured human DRG, CB2 agonists prevent capsaicin-induced Ca^2+^ influx indicating CB2 receptors are indeed present on cultured nociceptive primary afferent neurons (61). Future studies are needed to probe the specific subtypes of peripheral sensory afferents that contribute to the antinociceptive effects of CB2 receptor stimulation. Furthermore, future studies could examine whether CB2 is dynamically altered, induced or upregulated on certain primary afferent subtypes by specific pathologies. Such studies can be expected to provide further insight into particular pain conditions that may, or may not, benefit from CB2 receptor agonist therapy.

In the current study, the CB2 agonist LY2828360 robustly prevented the development of morphine tolerance in male CB2^f/f^ mice, and to a lesser extent in female CB2^f/f^ mice. To our knowledge, our study is the first demonstration of the tolerance sparing effects of a CB2 agonist in female mice. Mechanisms underlying differences between the tolerance sparing effects of LY2828360 in male and female mice in the present studies requires additional examination.

The present study is the first report to indicate that treatment with a CB2 agonist can reverse established morphine tolerance. The ability of a pharmacological intervention to *reverse* tolerance once established may prove critical in a clinical setting, as there are great individual differences regulating the development of tolerance in clinical populations (62). Once again, LY2828360 more robustly reversed established morphine tolerance in male versus female CB2^f/f^ mice. Although there have been many studies examining the existence of sex differences in opioid analgesia and tolerance ((63) for review), the current study failed to find any reliable differences in opioid analgesia or tolerance between male and female mice. The difference in LY2828360’s ability to prevent or reverse morphine tolerance is unlikely to be attributed to differences in the kinetics of the development or extent of morphine tolerance between male and female mice; in the absence of LY2828360, mice of both sexes developed tolerance to morphine’s ability to alleviate ddC-evoked mechanical and cold hypersensitivity at equivalent rates and to a similar extent. Further experiments are necessary to examine whether differences in CB2 receptor expression between male and female mice can be detected in regions of nociceptive circuitry controlling the development of morphine tolerance (such as primary afferent neurons (16)).

The present results build on previous work from our laboratory (22) indicating that administration of LY2828360 prior to or co-administered with morphine prevents the development of tolerance to morphine’s antinociceptive efficacy in a model of chemotherapy-induced peripheral neuropathy. Previous studies from other laboratories have suggested CB2 stimulation prevents the development of morphine tolerance by upregulating MOR mRNA levels in DRG and spinal cord (64), or by reducing tolerance produced by the release of proinflammatory cytokines via stimulation of CB2 receptors on microglia (65). The possibility also exists that LY2828360 may more effectively engage either of these pathways in male mice. Further experiments are required to determine whether CB2 receptor stimulation produces greater upregulations of MOR mRNA in males relative to female mice or suppresses opioid induced increases in proinflammatory cytokines more effectively in male versus female mice. However, the link between stimulation of MOR on microglial cells and the development of tolerance is in dispute (16, 66–68), as MOR mRNA appears absent in microglia (16). Recently, the critical role of MOR’s on peripheral sensory neurons in driving pro-nociceptive plasticity thought to underlie the development of tolerance to opiates has been demonstrated (16). The results of the current study support these findings, as the tolerance sparing and reversing effects of the CB2 agonist LY2828360 were completely absent in male and female mice lacking CB2 on peripheral sensory neurons. Future studies should examine the particular subtypes of sensory neurons (e.g., nociceptors vs. mechanoreceptors) which underlie the tolerance sparing effects of CB2 agonists on morphine antinociception, as well as the signaling mechanisms underlying these effects. Additionally, whether CB2 receptor stimulation decreases pro-nociceptive plasticity induced by morphine in peripheral sensory neurons via direct (e.g., CB2 stimulation directly decreasing firing rates of peripheral sensory neurons), or indirect mechanisms (e.g., CB2 stimulation on microglia decreasing the release of proinflammatory factors driving tolerance) is poorly understood. Whether pathological conditions are necessary to induce or alter CB2 expression on subsets of peripheral sensory neurons that, in turn, are necessary for the tolerance sparing effects of CB2 receptor stimulation requires further evaluation.

The present studies support the use of CB2 agonists in alleviating HIV-associated antiretroviral neuropathy and demonstrate, for the first time, that CB2 expression on peripheral sensory neurons are necessary for the antinociceptive efficacy of CB2 agonists. Our studies also demonstrate that the CB2 agonist LY2828360 prevents the development of antinociceptive tolerance to morphine and provides the first evidence that a CB2 receptor agonist *reverses* established tolerance to morphine antinociceptive efficacy in a neuropathic pain model. Furthermore, our studies have provided the first evidence that CB2 receptor expression on peripheral sensory neurons is necessary for the anti-allodynic efficacy of CB2 agonists as well as the ability of a CB2 agonist to prevent or reverse morphine tolerance.

## Materials and Methods

### Subjects

Adult (approximately 3 months old at the start of experiments) male and female C57Bl6/J mice were purchased from Jackson labs (Bar Harbor, ME). Adult (3-5 months old at start of experiments) CB2 knockout mice, advillin^Cre/+;^CB2^f/f^ and CB2^f/f^ mice were bred at Indiana University on a C57Bl6/j background. All procedures were approved by the Institutional Animal Care and Use Committee at Indiana University, Bloomington.

### Drugs

2’-3’-dideoxycitidine (ddC) was purchased from Sigma Aldrich (St. Louis, MO). AM1710 was synthesized by the lab of Alexandros Makriyannis (Northeastern University, Boston, MA). LY2828360 was synthesized by Sai Life Sciences Ltd. (Hyderabad, India) for use in all experiments with the exception of experiment 1. LY2828360 used in experiment 1 was provided by Eli Lilly (Indianapolis, IN) and used to validate results of LY2828360 synthesis from Sai Life Sciences Ltd. Morphine sulfate was provided by the NIDA Drug Supply Program (Bethesda, MD). ddC was dissolved in 0.9% sterile saline (Baxter Healthcare Corporation, Deerfield, IL). All other compounds were dissolved in a vehicle consisting of 3% DMSO, and the remaining 97% consisting of ethanol, emulphor, and 0.9% saline at a ratio of 1:1:18 respectively. All compounds were delivered at a volume of 5 ml/kg via intraperitoneal (i.p.) injection 30 minutes before behavioral testing occurred.

### Behavioral hypersensitivity testing

Paw withdrawal thresholds to mechanical stimulation were measured using an electronic von Frey anesthesiometer (IITC, Woodland Hills, CA) attached to a semiflexible plastic tip as described previously (14, 22, 69–71). Animals were placed on a mesh table under Plexiglas chambers and allowed to habituate to the environment for 30 minutes prior to the onset of behavioral testing. Mechanical stimulation was applied to the midplantar surface of the hind paw and the amount of force in grams necessary to provoke a withdrawal response was recorded in duplicate for each paw. Cold allodynia was assessed by applying a droplet of acetone to the midplantar surface of the hind paw and measuring the amount of time spent attending to the stimulated paw (i.e. licking, shaking, and lifting of the paw) in triplicate for each paw (14, 22, 69–71).

### General experimental procedures

ddC was administered 3 times per week for a total of 3 weeks at 25 mg/kg, i.p (cumulative dose: 225 mg/kg i.p. over 3 weeks) as described previously (18, 20). For all experiments, pharmacological testing began on day 14 after the initiation of ddC treatments when mechanical and cold allodynia were maximal and stable. Within-subjects ascending dose response curves were generated allowing four days to pass between each pharmacological treatment. In experiment 1, the development of ddC-induced mechanical and cold allodynia was measured three times per week (Monday, Wednesday, Friday). In all subsequent experiments, behavioral responsiveness during the development of ddC-induced pain was measured every 4 days. In the experiment utilizing CB2 KO mice, male and female CB2 KO mice were treated with a single injection of AM1710 or LY2828360. Then, 24 h later, the same subjects received a single i.p. injection of morphine. In chronic dosing experiments, mice received once daily injections of AM1710, LY2828360 or morphine beginning on day 14 of ddC treatment. The time course of acute drug treatments was determined on day one of chronic dosing. In experiments examining the ability of LY2828360 to prevent or reverse morphine tolerance, mice received once daily injections of morphine (10mg/kg, i.p.) or vehicle, in combination with LY2828360 (0.03 mg/kg or 0.1 mg/kg, i.p.) or vehicle for the co-administration dosing schedule. For the reversal dosing schedule, mice received once daily injections of morphine (10 mg/kg, i.p.) or vehicle for all 12 days of chronic dosing, in combination with vehicle for days 1-6, followed by LY2828360 (0.03 mg/kg or 0.1 mg/kg) or vehicle, on days 7-12 of administration. For all conditions, mice were tested every three days to evaluate the development of tolerance, and behavioral testing began 30 minutes after the injection of the pharmacological treatments. All experiments were conducted by experimenters blinded to treatment conditions and genotype. Sample sizes were determined using pilot data from our laboratory. Experiments were run using multiple batches of littermates with each treatment condition represented per batch of experiments. Animals were randomly assigned to groups using a random number generator. Baseline levels of mechanical and cold sensitivity were analyzed for each treatment group to ensure no differences existed at baseline.

### Tissue preparation for immunohistochemistry/methods for immunohistochemistry

Male and female CB2^f/f^ mice used for immunohistochemical experiments received ddC (25 mg/kg i.p./day 3x/week) for a total of 2 weeks and were sacrificed on day 14 following the onset of treatments to coincide with the time point used to initiate pharmacological treatments in all other experiments. Tissue was harvested approximately 48 hours after last ddC injection. Mice were deeply anesthetized with isoflurane and then transcardially perfused with 0.1% heparinized 0.1M phosphate-buffered saline (PBS) followed by ice cold 4% paraformaldehyde. Spleens, lumbar spinal cords, and brains were extracted and kept in the same fixative for 24 hours and then cryoprotected in 30% sucrose for 3 days prior to sectioning. Lumbar L3-L6 dorsal root ganglia (DRG) were extracted, kept in the same fixative for 4 hours and cryoprotected in 30% sucrose overnight prior to sectioning. Brains, spleens, and spinal cords were cryosectioned at 30 μm and maintained in an antifreeze solution (50% sucrose in ethylene glycol and 0.1M PBS) prior to immunostaining. Free floating sections were washed in 0.1M PBS, blocked with buffer consisting of 5% donkey serum and 0.1% Triton X-100 in 0.1M PBS, then incubated overnight in goat polyclonal antibody to GFP (AB0020-200, 1:1000, Sicgen, Cantenhede, Portugal) at 4° C. Sections were then incubated in donkey anti-goat AlexaFluor 594 (1:500, ThermoFisher, Waltham, MA) counterstained with DAPI (0.1 μg/mL) and treated with Sudan black B (0.2% weight/volume, Sigma Aldrich, St Louis, MO). DRG were cryosectioned at 12 μm, mounted on gelatin-subbed slides, and kept at −80° C prior to immunostaining. Slide-mounted immunostaining was carried out using identical procedures as free-floating immunostaining.

### Digital imaging and microscopy

Images were captured using a Leica DM6 B microscope equipped with a Leica DFC9000 GT camera and Leica Application Suite X software (Leica Microsystems, Wetzlar, Germany).

### RNA extraction and quantitative reverse transcription polymerase chain reaction (qRT-PCR)

Naïve mice were deeply anesthetized with isoflurane, and lumbar spinal cord, DRG, and spleen were dissected and flash frozen in isopentane and stored at −80 °C until use. Total RNA was purified using TRIzol^TM^ (Thermofisher, Waltham, MA) and RNeasy mini kit from lumbar spinal cord, spleen, and dorsal root ganglia according to manufacturer’s manual. Quantification of total mRNA was assessed by using a NanoDrop 2000C UV-Vis spectrophotometer at 260 nm. One-step RT-qPCR was performed using the Luna universal One-Step RT-qPCR kit (New England BioLabs) in a total volume of 20 µl and a template concentration of 50 ng/µl according to the manufacturer’s recommendations. Thermal cycle conditions were 55 °C for 10 minutes (RT step), 95 °C for 1 minute, followed by 40 cycles of 95 °C for 10 seconds and 60 °C for 1 minute. A melting curve analysis was performed at 95 °C for 15 seconds, 60 °C for 1 minute, 95°C for 15 seconds following every run to ensure a single amplified product for each reaction. All reactions were performed in duplicates in 96 well reaction plates (Thermofisher, Waltham, MA). Primer sequences are listed as follows:

1. Forward GAPDH/Reverse GAPDH: GGGAAGCTCACTGGCATGGC/GGTCCACCACCCTGTTGCT
2. Forward CB2/Reverse CB2: CTCGGTTACAGAAACAGAGGCTGATGTG/ TCTCTCTTCGAGGGAGTGAACTGAACG
3. Forward GFP/Reverse GFP: ACATGGTCCTGCTGGAGTTCGTGAC/CTCTTCGAGGGAGTGAACTGAACGG

For experiments evaluating impact of ddC and pharmacological treatments of cytokine/chemokine mRNA expression levels, RT-qPCR was performed using Taqman RNA-to-CT 1-step kit (Thermo fisher scientific, Waltham, MA) in a total volume of 10 µl and a template concentration of 50 ng/µl according to manufacturer’s recommendations. Thermal cycle conditions were 48 °C for 30 minutes (RT step), 95 °C for 10 minutes, followed 40 cycles of by 95 °C for 15 seconds, and 60C °C of 1 minute. A melting curve analysis was performed at 95 °C for 15 second, 60 °C for 1 minute, 95 °C for 15 seconds following every run to ensure a single amplified product for every reaction. Taqman gene expression assays used in the experiments were: GAPDH mm9999995-g1; CNR2 m02620087_s1; IL10 mm00439616_m1; IL-1beta mm00434228_m1; CXCL12 mm00445553_m1; CXCR4 mm01996749_s1; TNFα mm00443258_m1; CCL2 mm00441242_m1.

All reactions were performed in duplicate in 394 well reaction plates (Thermofisher Scientific, Waltham, MA). The mRNA expression levels were expressed as relative quantities to control and calculated by 2(-ΔΔCt) method where GAPDH is used as in house reference gene.

### Statistical analyses

Paw withdrawal thresholds and duration of time spent attending to the acetone-stimulated paw were averaged across paws for each modality (mechanical, cold) for each subject on each day to obtain a single data point per subject per observation interval. The time course of the development of ddC-induced neuropathic pain, as well as within-subjects dose-response curves, were analyzed by two-way repeated measures ANOVA followed by Tukey multiple comparisons tests post-hoc to examine differences between groups. Unpaired sample two-tailed t-tests were used for two group comparisons to analyze qPCR data. Statistical analyses an all figures were produced using Graphpad prism 7 (Graphpad prism software, San Diego, CA).

## Acknowledgements

This work was supported by National Institute on Drug Abuse Grants DA047858, DA041229 (A.G.H. and K.M.), and DA042584 (A.G.H.), National Cancer Institute Grant CA200417 (A.G.H.), an Indiana Addiction Grand Challenge Grant (A.G.H.), the Research and Education Component of the Advancing a Healthier Wisconsin Endowment at the Medical College of Wisconsin (C.J.H) and the Ministerio de Economía y Competitividad (SAF 2016-75959-R and SAF PID2019-108992RB-I00 to JR). L.M.C. was supported by National Institute on Drug Abuse T32 training grant DA024628 and the Harlan Scholars Research Program.

